# Are interphylum spiralian relationships resolvable?

**DOI:** 10.64898/2026.01.25.701568

**Authors:** Ana Serra Silva, Maximilian J. Telford

## Abstract

The phyla making up the major animal clade of Spiralia have been clear since the advent of molecular phylogenetics; the relationships between these spiralian phyla have not. The lack of consensus over the relationships between these important animal phyla might be a clue implying their emergence in an explosive radiation. Focussing on the five largest spiralian clades (Annelida, Brachiozoa, Mollusca, Nemertea and Platyhelminthes) and using two phylogenomic datasets, we have applied site-bootstrapping and taxon-jackknifing to explore this example of taxonomic instability. Analyses on the 105 possible rooted trees relating them showed that interphylum branches are very short. Preference for rooting Spiralia on Platyhelminthes is enhanced by a long-branch artefact. Most analyses on the 15 unrooted trees showed a preference for the same topology but the support for this tree over other solutions was not significant. We conclude that the spiralian phyla emerged in rapid succession resulting in a difficult to resolve radiation. The deep history we infer for Spiralia has wide ranging implications for our interpretation of Cambrian fossils and for the evolution of traits such as biomineralization, segmentation and larvae.

**Impact Statement:** Analyses of two independent phylogenomic datasets suggest an explosive radiation at the origin of Spiralia, with implications for understanding the group’s evolutionary history.

## Introduction

The group of animal phyla with members that undergo spiral cleavage in early embryogenesis were first shown to be closely related in the late 1990s using molecular sequence data (Halanych et al., 1995). The clade Spiralia unites the spirally cleaving Mollusca, Annelida, Nemertea, Platyhelminthes, Brachiozoa (including Brachiopoda and Phoronida) and Entoprocta, as well as Gastrotricha and Ectoprocta/Bryozoa. Spiralia forms the best part of the greater Lophotrochozoa (Halanych et al., 1995), the sister clade of Ecdyzosoa, and constitutes a significant part of animal diversity with close to 100,000 species of molluscs, 20,000 flatworm species, almost as many annelids, and several other smaller phyla. The sister group of Spiralia within Lophotrochozoa is the Gnathifera clade (Marlétaz et al., 2019; Vinther and Parry, 2019) which unites several phyla of jaw-bearing animals—Chaetognatha and the tiny Gnathostomulida, Rotifera and Micrognathozoa. Given the often-interchangeable use of the names Lophotrochozoa and Spiralia, we provide an abridged history of the uses of these names in the supplementary information.

Soon after the recognition of Lophotrochozoa (Aguinaldo et al., 1997; Halanych et al., 1995), it was found that the internal branches connecting its constituent phyla were short (Halanych, 2004), a phenomenon confirmed by the use of large phylogenomic datasets (Fig. 1). Several sources of systematic error seem to be at play in this clade, including lineage- and site-compositional heterogeneity (Nesnidal et al., 2010; Nesnidal et al., 2013), and branch-length heterogeneity (Nesnidal et al., 2010; Nesnidal et al., 2013; Struck et al., 2014). Attempts at mitigating these sources of error yielded a set of sometimes well-supported yet conflicting topologies (Laumer et al., 2019), or yielded topologies with very poorly supported internal branches (Kocot et al., 2017; Struck et al., 2014).

**Fig. 1.**
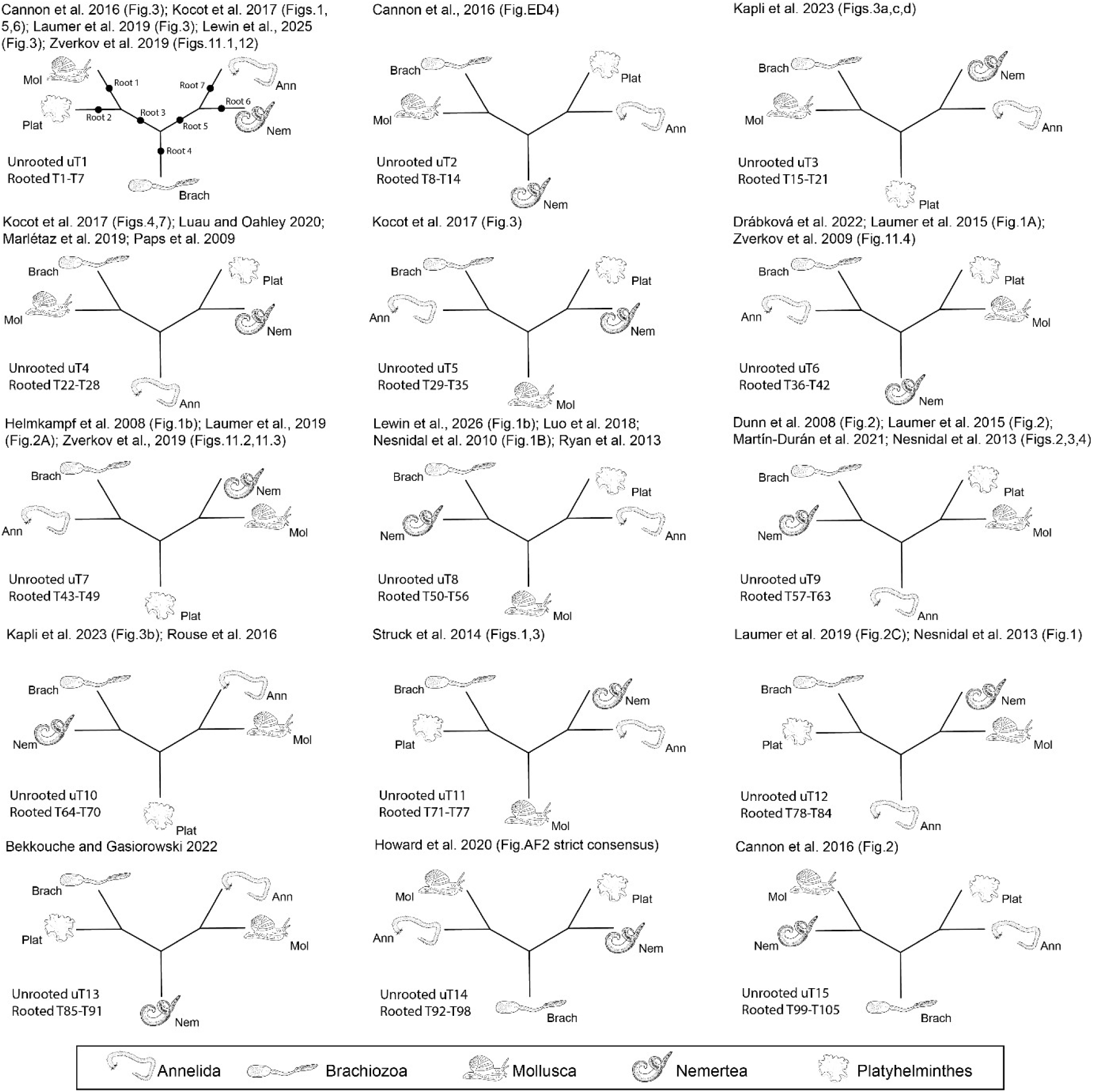
There are 15 unrooted trees linking the 5 major spiralian phyla and each has been supported in at least one publication (table S1). Tree numbering throughout this study refers to the trees in this figure. For all 105 rooted trees, the rooting schemes follow that depicted in unrooted T1.

As evidence of the lack of consensus, multiple, often contradictory, subclades have been proposed and named based on the many relationships inferred with molecular phylogenetics: Parenchymia (Platyhelminthes/Nemertea/Annelida) (Marlétaz et al., 2019); Rouphozoa (Platyhelminthes/Gastrotricha) (Struck et al., 2014); Tetraneuralia (Mollusca/Entoprocta) (Wanninger, 2009); Platyzoa (Platyhelminthes/Gnathifera/Gastrotricha) (Cavalier-Smith, 1998); Trochozoa (‘Clade C’ in Dunn et al. (2008): Mollusca/Annelida/Nemertea/Brachiopoda/Phoronida) (Giribet et al., 2000); Eutrochozoa (Mollusca, Annelida, Nemertea) (Ghiselin, 1988); Vermizoa (Annelida/Nemertea) (Cavalier-Smith, 1998); Conchozoa (Mollusca/Brachiopoda/Phoronida) (Cavalier-Smith, 1998); Platytrochozoa (Rhouphozoa/Trochozoa) (Struck et al., 2014); Kryptrochozoa (‘Clade A’ in Dunn et al. (2008): Nemertea/Brachiopoda/Phoronida) (Giribet et al., 2009); ‘Clade B’ (Nemertea/Brachiopoda/Phoronida/Annelida) (Dunn et al., 2008); Neotrochozoa (Mollusca/Annelida) (Peterson and Eernisse, 2001); Plathelminthomorpha (Platyhelmithes/Gnathostomulida) (Ax, 1984); Monokonta (Gastrotricha/Gnathostomulida) (Cavalier-Smith, 1998); Brachiozoa (=Phoronozoa, Brachiopoda/Phoronida) (Cavalier-Smith, 1998; Zrzavý et al., 1998); etc.

The confusion over the relationships between spiralian taxa is also obvious in the lack of convincing morphological or embryological synapomorphies linking subsets of phyla. Schleip (1929), in his paper coining the name Spiralia (for what he explicitly considered a non-monophyletic assemblage), illuminates the problem we want to address: “The spiral type of cleavage is found in a number of animal groups, whose organisation in the developed state [i.e. adults] is quite different.” While it is now widely accepted that the spiralians *are* united by unique features of their early development (which in many includes a ciliated larval stage) the relationships *between* the spiralian phyla seem to have been obscured by their very different adult forms (Eernisse et al., 1992; Vinther and Parry, 2019). Synapomorphies for different possible collections of spiralian phyla are of enormous evolutionary interest: are annelids and molluscs (and perhaps brachiopods; Gutmann et al., 1978; Temereva and Malakhov, 2011) united by a form of segmentation? Are the chaetae of annelids and brachiopods and/or the shells of brachiopods and molluscs homologous (Liang et al., 2025; Schiemann et al., 2017)?

This lack of synapomorphies and the breadth of phylogenomic studies finding strong support for incongruent spiralian interphylum relationships raises the possibility of a multifurcation at the origin of this clade. Despite the now widespread acceptance of non-tree-like processes (Ishii et al., 2026; Song et al., 2023) and non-bifurcating speciation (Helmkampf et al., 2025; Parins-Fukuchi and Saulsbury, 2025), proof of multifurcation events remains elusive, even for comparatively recent radiations (Scherz et al., 2022; Suh, 2016). Proving a hard polytomy at the base of Spiralia, however, requires dismissing the binary topologies that might relate spiralian phyla. This is not trivial given the expected loss of phylogenetic signal (Mossel and Steel, 2005; Penny et al., 2001; White et al., 2007) resulting from the group’s age (>543.0 million years according to Carlisle et al. 2024) and the various known sources of systematic error (Nesnidal et al., 2010; Nesnidal et al., 2013; Struck et al., 2014).

We have used two recent phylogenetic datasets to ask whether there is convincing support for any clades uniting more than one spiralian phylum. We have reduced our taxon sampling to five principal phyla: Annelida, Mollusca, Platyhelminthes, Nemertea and the Brachiopoda (in which we include its sister phylum Phoronida and hereafter use the superphyletic Brachiozoa clade name) and the outgroups Gnathifera, Ecdysozoa, Chordata, Ambulacraria and Cnidaria. These five monophyletic spiralian groups can be linked by 15 unrooted and 105 rooted trees. This simplification allows us to focus on direct comparisons of the support for each tree using different datasets, different models of evolution and different samples of taxa.

To characterise the level of uncertainty in interphylum spiralian relationships, we started by doing a literature review on previously inferred trees. Support for possible 5-taxon rooted and unrooted topologies was then compared under site-homogeneous and site-heterogeneous models, with both fixed and randomised taxon sampling. Lastly, we used simulations to identify artefactual signals in both the rooted and unrooted trees.

## Results

### Literature review shows historical support for all 15 possible unrooted spiralian trees

A non-exhaustive review of studies dealing with the phylogeny of Lophotrochozoa, focussing on studies that sampled all five spiralian target clades, showed that trees consistent with all 15 possible unrooted spiralian topologies have been inferred by at least one publication (Bekkouche and Gąsiorowski, 2022; Cannon et al., 2016; Dunn et al., 2008; Helmkampf et al., 2008; Howard et al., 2020; Kocot et al., 2017; Lau and Oakley, 2021; Laumer et al., 2015; Laumer et al., 2019; Luo et al., 2018; Marlétaz et al., 2019; Martín-Durán et al., 2021; Nesnidal et al., 2010; Nesnidal et al., 2013; Paps et al., 2009; Rouse et al., 2016; Ryan et al., 2013; Struck et al., 2014), see figure 1 and data D1. What these efforts reveal most strikingly is a lack of consensus on spiralian relationships. Even the most recent studies contradict each other, with some studies showing support for multiple topologies (Drábková et al., 2022; Kapli et al., 2023; Lewin et al., 2025; Lewin et al., 2026; Zverkov et al., 2019).

### Whatever the true topology, clades grouping phyla are linked by extremely short branches

The historical lack of clear resolution within Spiralia suggests that the branches separating clades may be short. We have used two recent phylogenomic data matrices (Marlétaz et al. (2019) and Serra Silva et al. (2025), MARL and SERR hereon) to explore the support for different trees linking the five principal spiralian phyla.

We used IQ-Tree v.2.3.5 (Minh et al., 2020) to compute log-likelihoods and to optimise branch lengths for each of the 105 rooted trees using both the site-homogeneous LG+F+G4 model and dataset-specific, site-heterogeneous, 128-category empirical distribution mixture (EDM, Schrempf et al., 2020) models. Intraphylum relationships were fixed using internally unconstrained maximum likelihood (ML) tree inferences (LG+F+G4) on both matrices. For both datasets, and on all 105 trees, we measured all internal branches linking our five phyla. We compared these lengths to the branch leading from Urbilateria to the deuterostomes, recently shown to be hard to resolve and, if real, very short (Kapli et al., 2021; Serra Silva et al., 2025). All comparisons between branch lengths are restricted to fixed dataset+model pairs.

For both MARL and SERR datasets and regardless of model, almost all internal interphylum spiralian branches are shorter than the deuterostome branch (percentage of shorter branches: SERR LG = 92.38%, SERR EDM = 90.48%; MARL LG = 90.48%, MARL EDM = 94.92%; Fig. 2 and table S1). The exceptions (where the internal branch is approximately equal to or slightly longer than the deuterostome branch) are found on trees where the root separates either the very long-branched Platyhelminthes from the four other spiralian phyla or where Nemertea and Platyhelminthes are sister taxa. These two conditions correspond to the majority of the trees that are highest-ranked under the site-homogeneous LG+F+G4 model and may be the result of a long-branch attraction (LBA) artefact (see later). The short branches are reflected in the low (<50%) quartet scores recovered from discordance-aware Astral-III v.5.1.3 (Zhang et al., 2017) summary trees of each dataset (see below and Fig. S1). We also measured the length of branches leading to each of the five spiralian phyla, measured from its most recent common ancestor with another phylum/phyla. These were longer than the median deuterostome branch, supporting the idea that the clades considered as phyla are defined by long stem-lineages. The branch leading to Platyhelminthes was always much longer than the other phylum branches; followed by those leading to Mollusca and Annelida.

**Fig. 2.**
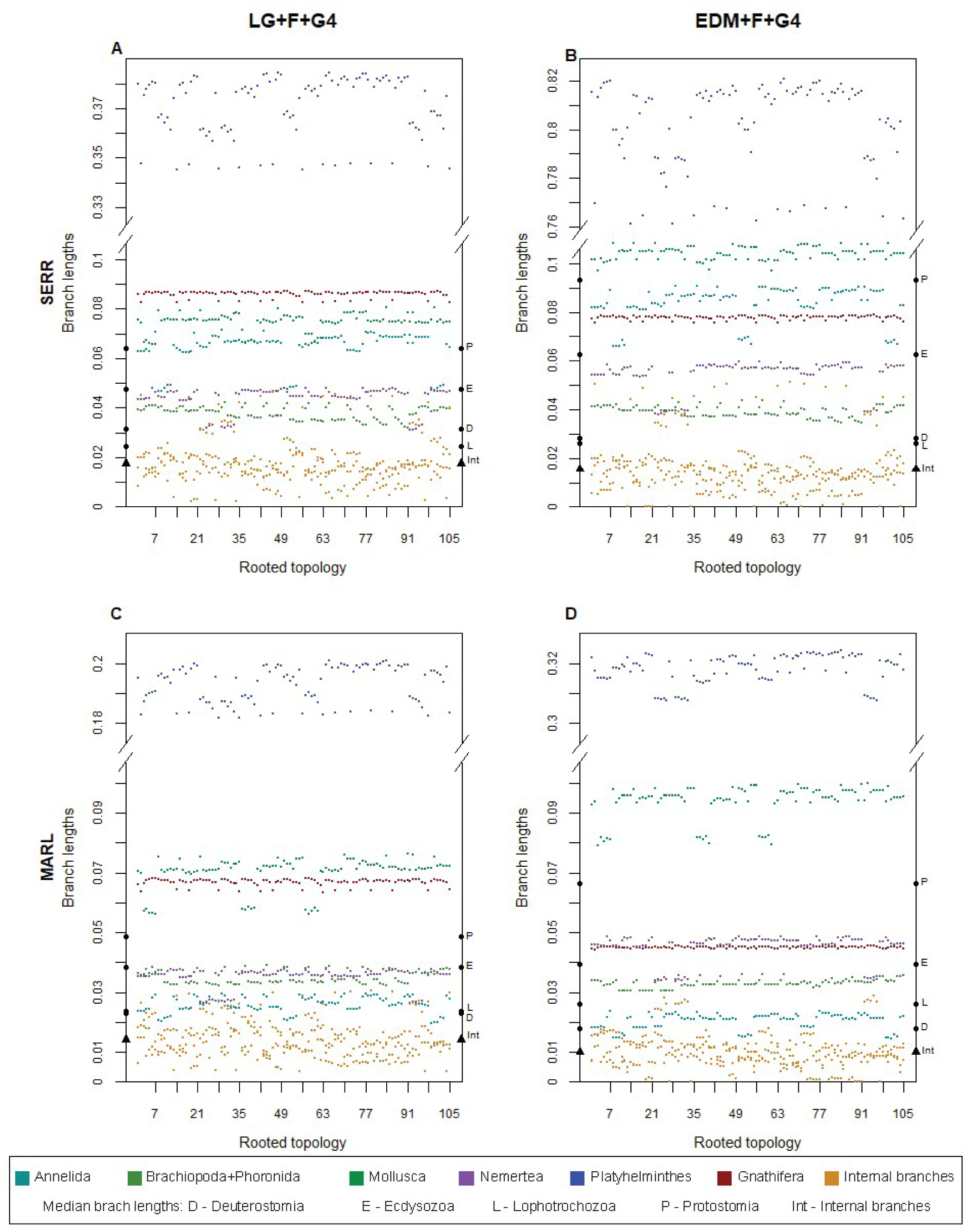
Branches linking spiralian phyla are shorter than both the short, disputed deuterostome branch (Kapli et al., 2021; Serra Silva et al., 2025) and the short branch leading to Lophotrochozoa (Telford et al., 2015). **A** and **B** are data from SERR. **C** and **D** are data from MARL. Compared to the site-homogeneous model (**A**, **C**), internal branches (orange) become shorter under the site-heterogeneous model while the long branches leading to Platyhelminthes (blue) and Mollusca (dark green) become even longer (**B**, **D**). Circles on the y-axis indicate the median branch lengths of Deuterostomia (D), Ecdysozoa (E), Lophotrochozoa (L) and Protostomia (P). Triangles correspond to the median spiralian internal branches (Int), values given in table S1. Scales are identical on either side of the y-axis breaks.

While not our primary interest, we note that the median length of the branch leading to Lophotrochozoa was similar to the short Deuterostome branch (Fig. 2 and table S1). This might explain the initial difficulties placing the chaetognaths within the protostomes with a confusion over whether they were ecdysozoans, lophotrochozoans or neither (Marlétaz et al., 2019; Telford et al., 2015).

With the site-heterogeneous EDM model, most internal branches become even shorter, while some of the longer branches leading to phyla (Platyhelminthes, Mollusca and Nemertea) become longer. We interpret this as site-heterogeneous models reinterpreting some characters shared by faster evolving phyla as having changed convergently (changing twice on external branches) rather than representing synapomorphies (single changes) on internal branches.

### Discordance-aware analyses find no support for any spiralian clades

Thus far, our likelihood-scoring analyses assumed that all loci in each dataset support the same topology (we have used an unpartitioned concatenated matrix). This assumption is often unmet due to gene-tree/species-tree incongruence, and this is especially likely when internal branches are short. To explore spiralian relationships in a gene-tree discordance-aware framework, we inferred two Astral-III summary trees, one each for SERR and MARL datasets, using individual locus ML gene-trees inferred under the site-homogeneous LG+F+G4 model in IQ-Tree. We did not attempt to identify whether gene-tree discordance was due to incomplete lineage sorting or reticulation events as these separate phenomena can yield the same gene-tree/species-tree discordance patterns (reviewed in Hibbins and Hahn, 2022), and we assumed binary trees for all topology scoring tests below.

Both supertrees (Fig. S1) recover monophyletic spiralian phyla, except for Annelida in the MARL supertree. This is consistent with previous work on the phylum, where *Owenia* is often found to be taxonomically unstable (Laumer et al., 2015; Rousset et al., 2007; Struck et al., 2015; Struck et al., 2008; Weigert et al., 2016). In both SERR and MARL Astral supertrees, the only internal spiralian branch to achieve quartet scores (QS) > 50% was the branch separating Platyhelminthes+outgroups from the remaining spiralian phyla. Platyhelminthes are recovered within a paraphyletic Gnathifera, which, in the context of the long Platyhelminthes and Gnathifera branches (Fig. 2), is likely to be an instance of LBA.

None of the other interphylum branches garner more than 50% QS, suggesting that we cannot distinguish between any of the various quartets involving Annelida, Mollusca, Nemertea and Brachioza. The high level of discordance seen across both datasets is consistent with the extremely short interphylum branches described above and is a pattern that has been recovered for multiple other (albeit considerably shallower) ‘hard to resolve’ clades such as treefrogs (Chan et al., 2020), birds (Sayyari and Mirarab, 2016) and lizards (Streicher et al., 2016).

Restricting the analyses to the gene-trees in which all phyla were monophyletic increased scores for the branches leading to phyla (not surprisingly) but not for interphylum branches. Restricting our analyses to the shortest gnathiferans and platyhelminths (measured from rooted T1) yielded similar topologies and quartet scores to those discussed above.

### Little support for specific topologies revealed by low bootstrap support and model dependency

We next asked whether our two datasets give consistent support for one or a few of the 105 possible rooted topologies linking the five clades (Annelida, Brachiozoa, Mollusca, Nemertea and Platyhelminthes) by comparing their relative ranks.

With the four sets of 105 likelihood-scored trees used for the branch length comparisons (i.e., LG and EDM for MARL and SERR), we used Kishino et al.’s (1990) resampling estimated log-likelihood (RELL) approach to generate large pseudo-site-bootstrap samples (10,000 replicates). We asked whether there were significant differences in support between topologies. We used the non-parametric Kruskal-Wallis test (KW, H-test) to ask whether the mean rank of the pseudo-bootstrap log-likelihoods of any one of the 105 rooted trees is significantly different from the others. This non-parametric test allows for comparisons between the single and multi-alignment (randomised taxon-jackknifing and simulations) analyses. Approximately unbiased (AU (Shimodaira, 2002)) tests with IQ-Tree yielded comparable results to our RELL+KW approach. The best-ranked topologies across pseudo-bootstrap replicates, ordered by median rank, are shown in figure 3 (see table S2 for all ranks).

**Fig. 3.**
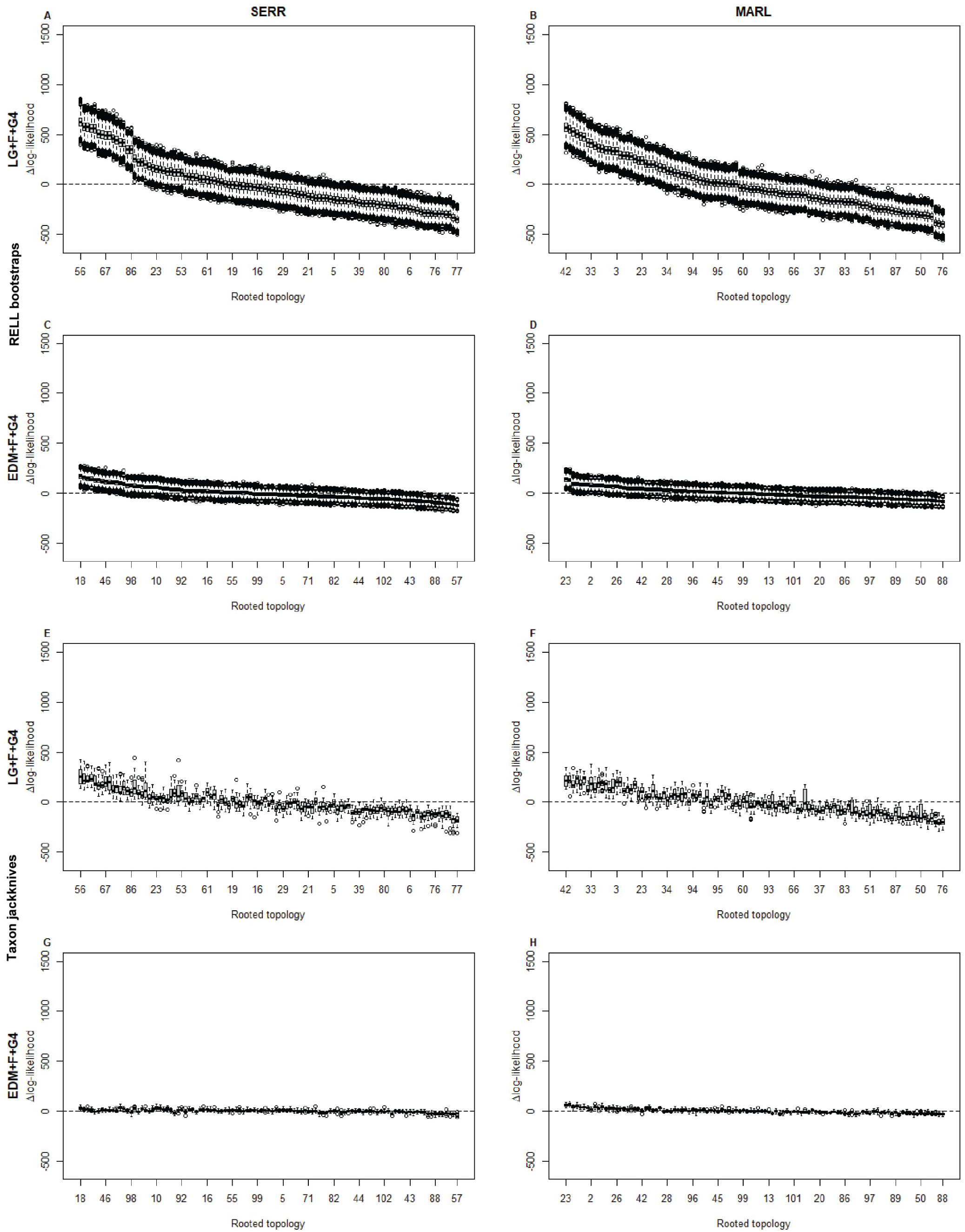
Relative median RELL pseudo-bootstraps (**A**–**D**) and randomised taxon-jackknives (**E**–**H**) log-likelihood ranks for all rooted trees, see tables S3 and S5 for all ranks. Highest- and lowest-ranked topologies labelled for all analyses, along with every seventh topology from highest- to lowest-ranked. Topologies ranked by their difference in log-likelihood from the mean log-likelihood of each RELL or jackknife distribution. The RELL analyses of both datasets under the site-homogeneous LG+F+G4 model (**A**, **B**) occupy the widest ranges of log-likelihoods, which is consistent with the statistically significant differences found by the KW test. The RELL analyses under the site-heterogeneous EDM+F+G4 model (**C**, **D**) occupy a much narrower range of log-likelihoods. In practical terms, the wider the log-likelihood range, the more dissimilar scores are and the more we would expect to recover significantly different scores between topologies. The jackknife analyses show a similar trend, wider range of median log-likelihood scores under LG+F+G4 (**E**, **F**) than under EDM+F+G4 (**G**, **H**); however, the log-likelihood ranges of the jackknives under LG+F+G4 are very similar to that occupied by RELL under EDM+F+G4, consistent with the lack of statistically significant differences between topologies in the jackknife analyses.

For the SERR data scored under the site-homogeneous model, the tree with the highest likelihood was unrooted T8 (uT8, for numbering see Fig. 1) rooted between Platyhelminthes and the other phyla. This tree was recovered as the highest-scoring across 63.53% of pseudo-bootstrap replicates (table S3). The 14 next highest scoring trees are the remaining 14 topologies also rooted on Platyhelminthes (table S2).

The KW test did find significant differences between topologies (p-value < 2.00 x 10^-16^), meaning that the 105 fixed-taxon sampling trees do not all have equivalent rank—some are consistently lower ranked than others. A *post hoc* Welch’s t-test showed that the 15 best trees (i.e. all those rooted on Platyhelminthes) are not significantly different from each other. Plotting the relative log-likelihoods of the pseudo-bootstraps of all 105 trees, we can see the 15 best trees have a higher likelihood than the worst but that median log-likelihoods of the best trees are very similar to one another and statistically indistinguishable (Fig. 3).

We found similar results using the site heterogeneous model—the 12 best supported trees are rooted on Platyhelminthes (Fig. 3 and table S2). The remaining three topologies rooted on Platyhelminthes also ranked highly (17, 20 and 27 of 105). The first ranked tree and the subsequent rank order of topologies is different to that of the LG+F+G4 analysis, uT3 rooted on Platyhelminthes is the highest ranked tree under EDM (Fig. 3). The KW test was non-significant (p-value = 1.00), meaning that there is no significantly preferred (most highly ranked) tree.

Analyses using MARL yielded qualitatively similar results, although the ranking of best supported trees is different. The site-homogeneous LG model still favours rooting on Platyhelminthes (ten of the 15 highest-scoring topologies were rooted on Platyhelminthes). The highest-scored tree corresponds to uT6 rooted on Platyhelminthes (highest-ranking in 50.33% of pseudo-bootstrap replicates: Fig. 3 and table S3). The KW test was significant (p-value < 2.00 x 10^-16^) but the *post hoc* t-test showed that the highest-ranked tree is not significantly different from the next ten highest-scoring topologies, all but two of which are rooted on Platyhelminthes.

When we apply the site-heterogeneous EDM model to the MARL data, in contrast, five of the 15 highest-scoring topologies are rooted on Mollusca (table S2). The highest ranked tree matches the topology reported in the original publication (uT4 rooted on molluscs). Only two of the top 15 are rooted on Platyhelminthes (ranks 8 and 10). As with the SERR data, the KW test was non-significant (p-value = 1.00).

These fixed taxon-sampling analyses suggest a significant preference for rooting on the branch leading to Platyhelminthes under the site-homogeneous LG+F+G4 model, while the analyses under the site-heterogeneous EDM model suggest non-significant dataset-specific preferences for rooting on Platyhelminthes (SERR) or Mollusca (MARL). The rooting on Mollusca seems less likely to be an artefact (see simulations). There is, in contrast, no clear result regarding the internal relationships of the five spiralian phyla.

### Taxon sampling reveals low support for any specific topology

To complement our site-bootstrapping approach, we explored the stability of support for different topologies under varying taxon sampling. We used randomised taxon-jackknifing to generate multiple subsampled replicates of both SERR and MARL datasets. For both datasets, we generated 20 jackknifed alignments with a total of 31 (SERR) or 30 taxa (MARL). We included a constant set of non-lophotrochozoan outgroups and from each spiralian phylum and from Gnathifera we sampled three taxa at random. We repeated likelihood-scoring analyses under the site-homogeneous (LG+F+G4) and site-heterogeneous (EDM+F+G4) models for all replicates under all 105 rooted spiralian topologies.

Under the site-homogeneous LG model, and based on median log-likelihood ranks, the six best-scoring trees in the SERR jackknives were rooted on Platyhelminthes (Fig. 2 tables S4–5), with both T18 and T56 recovered as the highest-ranked topology in 30% of replicates. For the MARL data, the two best-scoring trees also had this root and T18 was recovered as the highest-ranked in 25% of jackknife replicates. These rooting preferences were non-significant in both datasets (KW p-value = 1.00).

Under the site-heterogeneous EDM model, both MARL and SERR datasets narrowly favour a root on the branch separating Mollusca+Brachiozoa from the other spiralian phyla (T24 highest-ranked topology in 55% of MARL and 40% of SERR replicates) but the KW tests found no significant differences (KW p-value = 1.00).

### Simulations suggest that rooting on Platyhelminthes gets support from a long-branch attraction artefact

The most consistent result from our topology-scoring experiments is that there is a preference for rooting on the branch leading to Platyhelminthes or to Mollusca. Support for a platyhelminth root is reduced using a substitution model that accounts for between-site compositional constraints (the EDM model) suggesting that some of the support for this root might derive from an LBA artifact.

We have used a simulation approach using a known, unresolved topology (i.e. no root preferred) to explore whether the observed preference for rooting on Platyhelminthes, Mollusca, or Mollusca+Brachiozoa can emerge as artefacts through unaccounted for site-specific compositional constraints.

We first estimated model parameters using dataset-specific site-heterogeneous models for both MARL and SERR using an unresolved Spiralia tree (i.e. one in which the five phyla emerge as a polytomy). Using these parameters estimated under the site-heterogeneous EDM+G4 model, we simulated two corresponding sets of 100 alignments. Because the simulations were based on an unresolved tree, the support for the 105 different rooted trees that could relate the five phyla should, in the absence of artefact, be approximately equal.

For both datasets, we first scored the 105 rooted topologies under the correctly specified EDM model(s). As expected, we found no significantly preferred topology using the KW test (p-value = 1.00). In MARL, the two top-ranked topologies rooted on the branch leading to Platyhelminthes, while for SERR the top-ranked tree rooted on Platyhelminthes and the second-best on Mollusca (table S6). The preference for these over the others is tiny and not statistically significant.

Using a misspecified site-homogeneous LG model, we found a strong preference for rooting on Platyhelminthes (Fig. 4A–B). The 15 different topologies that have this root position outscored all other topologies and all other root positions (KW p-value < 2.00 x 10^-16^). For the MARL data, 84% of simulated alignments specifically recovered the uT5 topology rooted on Platyhelminthes as the highest ranked topology. While for the SERR data, 98% of simulated alignments recovered uT9 rooted on Platyhelminthes as the best-ranked tree, mirroring results from empirical data using the LG model.

**Fig. 4.**
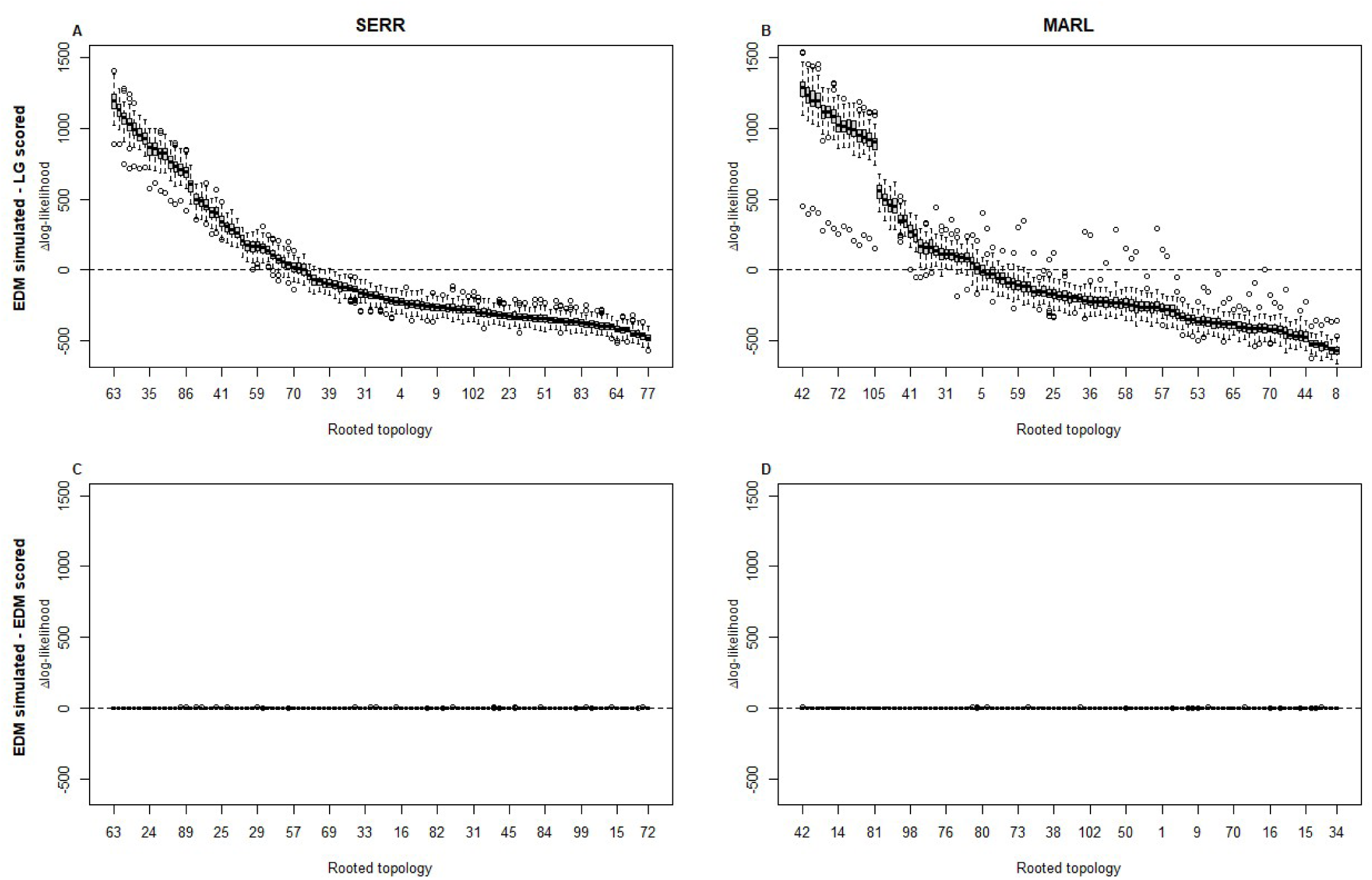
With simulated data inadequate models incorrectly support a platyhelminth root. Using data simulated on an unresolved Spiralia tree, we ranked rooted spiralian topologies by difference to mean log-likelihood score. Highest- and lowest-ranked topologies are labelled for all analyses, along with every seventh-topology from highest- to lowest-ranked. For datasets simulated under EDM (complex) but scored under LG (simple) the 15 topologies with Spiralia rooted on Platyhelminthes are highest ranked (**A**, **B**). The plots for the ‘correctly’ specified models (**C**, **D**) show the expected approximately equal log-likelihood scores for all 105 rooted topologies. See table S7 for complete topology rankings.

These simulations suggest that the strong preference for rooting on Platyhelminthes observed with empirical data is at least enhanced by unaccounted for site-compositional heterogeneity in the presence of unequal branch lengths.

### Removing the root does not unequivocally resolve relationships between spiralian phyla

Our simulations showed that preference for rooting Spiralia on Platyhelminthes is at least enhanced by a long-branch artefact exacerbated by site-compositional constraints. To circumvent the influence of the root, we removed the outgroups to Spiralia and asked whether any of the 15 unrooted, ingroup-only trees are significantly supported over the others. We repeated the fixed-taxon sampling, randomised taxon-jackknifing, and simulation analyses using the MARL and SERR datasets, under both site-homogeneous (LG+F+G4) and - heterogeneous (EDM+F+G4) models.

### Scoring 15 unrooted topologies using both datasets and with different models

All fixed-taxon topology-scoring analyses on the set of 15 unrooted topologies recovered the same highest-scored tree (uT4 in Fig. 1, table S8), with varying degrees of statistical significance. Under the site-homogeneous model, we find significant differences in rank between unrooted topologies (SERR KW p-value = 1.10 x 10^-4^; MARL KW p-value < 2.20 x 10^-16^). For the SERR data, comparing the highest-scoring topology to the other 14 with a *post hoc* Welch’s t-test showed that the highest-scored topology (uT4) was not significantly different from the topologies ranked two to five (uT2, uT14, uT8 and uT5) but was significantly preferred over the remaining ten topologies. With MARL, the *post hoc* t-tests found the highest-scored topology to be significantly different from all but the next highest scoring topology (uT5). Under the site-heterogeneous EDM, uT4 was not significantly better supported than any alternative (KW: SERR p-value = 1.00; MARL p-value = 0.80).

Pseudo-bootstrap rankings (Fig. 5 table S9), show a similar pattern to rooted analyses, with log-likelihoods of LG-scored topologies occupying a wider range of values than the EDM-scored topologies; however, unlike the analyses on the rooted topologies, the majority of pseudo-bootstrap replicates support the same unrooted topology, uT4 (SERR+LG = 81%, MARL+LG = 60%, SERR+EDM = 98%, MARL+EDM = 96%).

**Fig. 5.**
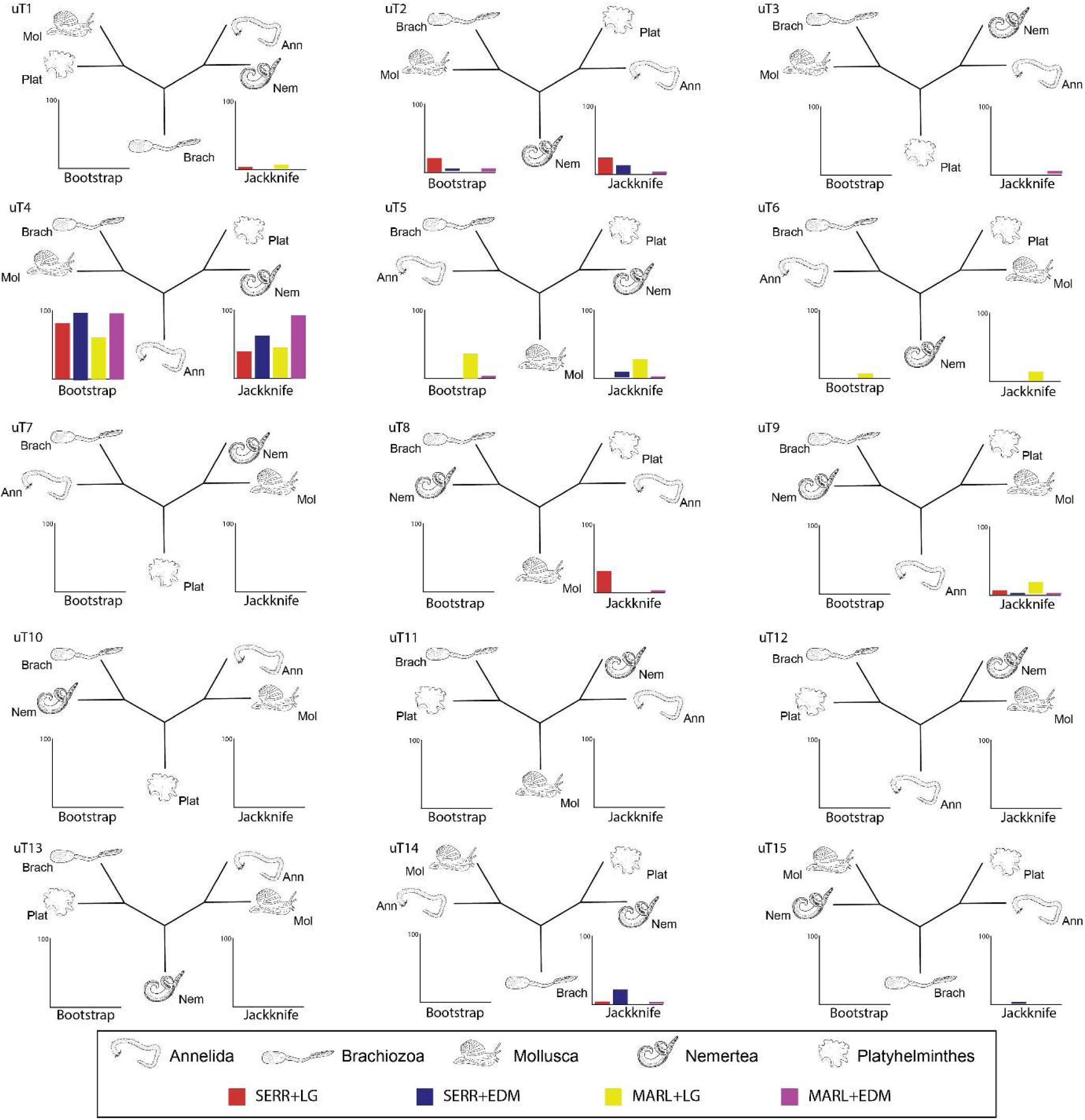
Most fixed-taxon RELL bootstrap and randomised taxon-jackknife replicates, under both site-homogeneous (LG) and site-heterogeneous (EDM) models, recover unrooted T4 as the best-scoring topology. Heights of bars correspond to the number of replicates where each topology had the highest log-likelihood score.

As with the analyses on the rooted topologies, when using a better fitting site-heterogeneous model (table S9) the 15 unrooted topologies are statistically indistinguishable. When using a site-homogeneous model, the highest-scored topology (uT4) is significantly better than some but not all alternatives.

### Taxon jackknifing on 15 unrooted trees shows species dependence

The taxon-jackknifing analyses on the 15 unrooted spiralian trees again showed uT4 was the highest-ranked topology in many but not all analyses (Fig. 5, table S10): uT4 had a slightly lower median log-likelihood than uT2 in the SERR+LG analyses (Fig. S2). The differences between topologies were not significant under either model for either dataset (KW p-value = 1.00). Although uT4 is most often found to be the best-supported tree, this depends on taxon sampling suggesting the support is not overwhelming. These results agree with the rooted taxon-jackknifing analyses, which found no *significantly* better supported in-group topology.

### Simulations using unresolved Spiralia only-tree

The simulations on the trees including outgroups suggested a site-composition bias in the data, with all analyses using a misspecified site-homogeneous model significantly supporting a root on Platyhelminthes. To test for an equivalent pattern of artefactual support in the Spiralia-only topologies, we repeated our simulations with an unresolved 5-taxon tree. As expected, with an adequate model we found no significant differences between the 15 unrooted topologies for either dataset (KW p-value = 1.00).

Using the misspecified LG model, the SERR-simulated data recovers no significant differences between the 15 5-taxon topologies (KW p-value = 1.00), although trees uT2, uT8, uT15, uT4, uT5, uT14 show the highest log-likelihood scores (Fig. 6 and table S11). For the MARL data, the misspecified LG analyses do show significant differences between topologies (KW p-value = 3.00 x 10^-6^). *Post hoc* t-tests find that the highest scoring topology (uT6) does not significantly differ from topologies uT1, uT4, uT5, uT9 and uT14, but uT6 is significantly different from the nine lowest scoring topologies.

**Fig. 6.**
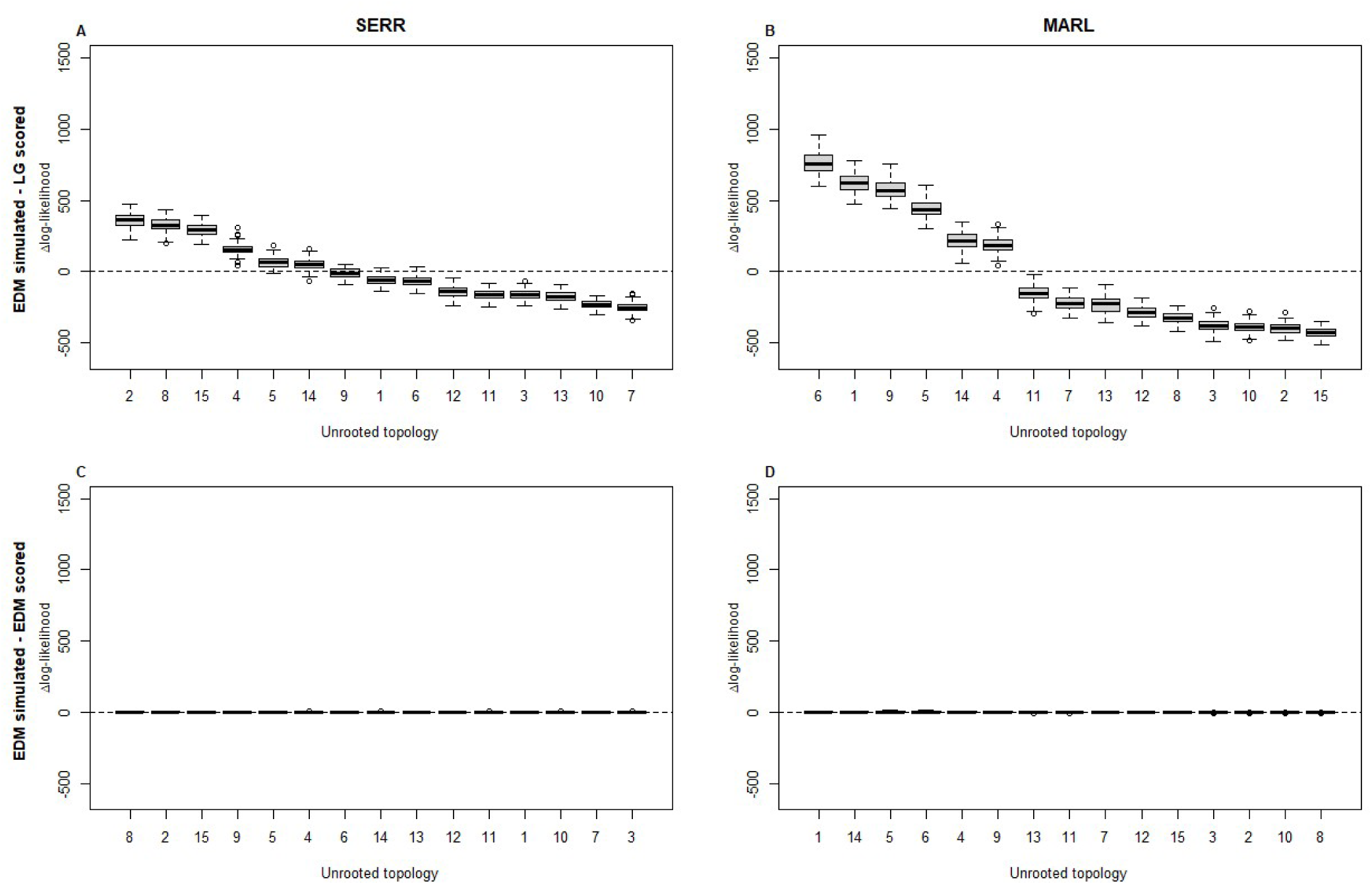
With simulated data, misspecified models recover unrooted T4 within the set of highest-scoring topologies. Using data simulated on an unresolved Spiralia tree we ranked unrooted spiralian topologies by difference to mean log-likelihood score. For datasets simulated under EDM (complex) but scored under LG (simple), three topologies (uT4, uT5 and uT14) are within the highest-ranked topologies for both empirical datasets (**A**, **B**). The plots for the ‘correctly’ specified models (**C**, **D**) show the expected approximately equal log-likelihood scores for all 15 unrooted topologies.

Inadequate models can, in short, result in limited artefactual support for a subset of the 15 unrooted topologies among the more favoured of which is the most frequently supported uT4 tree suggesting some of its support may be artefactual. Trees uT4, uT5 and uT14 are in the top 6 in both cases, and all contain the Nemertea+Platyhelminthes clade, suggesting that this grouping derives some support from systematic errors.

## Discussion

The relationships between the spiralian phyla have proved hard to pin down despite large datasets. Here, we have shown that the best-supported position of the root of the Spiralia is strongly affected by an LBA artefact between Platyhelminthes and the outgroup. We have used randomised-taxon jackknifing to show that the preferred topology can change according to the taxa used, suggesting a weak signal and this is also implied by the low quartet scores seen in the discordance-aware Astral analyses (Fig. S1) and in the very short internal branch lengths.

When we make efforts to tackle some of these problems, we observe most support for the (unrooted) topology uT4—((Mollusca,Brachiozoa),((Annelida),(Platyhelminthes, Nemertea))), which is also congruent with the topology inferred by Marlétaz et al. (2019). The support for this best supported topology is relatively weak, however, and, in the context of the very short internodes and evidence for other instances of error, makes us wary of declaring this to be *the* correct tree. Our simulation study further suggests that the relationship between platyhelminths and nemerteans seen in this tree receives at least some support from systematic error, moreover the correct position of the root on this tree is unknown.

Our experiments, using both empirical and simulated data, tackled only across-site compositional-heterogeneity under stationary amino acid frequencies and, as such, other models accounting for across-lineage compositional heterogeneity (e.g., CAT-BP (Blanquart and Lartillot, 2008), GHOST (Crotty et al., 2020), NDCH2 (Foster, 2022)); both across-site and across-lineage compositional heterogeneity (e.g., GFmix (McCarthy et al., 2025; Muñoz-Gómez et al., 2022)); non-stationary models (e.g., CAT-GTR (Lartillot and Philippe, 2004), of which the 128-EDM model is an approximation); or even multi-tree histories (e.g., MAST (Wong et al., 2024)) might yield different results. Even models like the CAT-GTR and CAT-BP, have not yielded cross-study support for *one* specific spiralian topology (Drábková et al., 2022; Nesnidal et al., 2010; Nesnidal et al., 2013). This lack of consensus, even with complex models, makes it hard to predict how models other than those used in this study (i.e. LG and EDM) might change the inferred/supported spiralian topologies. We might expect that artefactual support for rooting Spiralia on the branch to Platyhelminthes would be reduced with models accounting for lineage and/or site-compositional heterogeneity (but see Nesnidal et al., 2010) and that, congruent with our hard-polytomy-like interphylum quartet scores, multi-tree models would support multiple topologies across loci.

While our randomised taxon-jackknifing analyses show strong taxon-sampling effects on the best-supported topology (Figs. 3E-H and S2), our efforts to enable an exhaustive exploration of tree space (and to limit computational burden) by restricting our analyses to the five largest spiralian phyla mean that our findings are not applicable to the broader Spiralia—which includes Cycliophora, Ectoprocta/Bryozoa, Entoprocta, Gastrotricha and the extremely simplified endoparasites Orthonectida and Dicyemida (Drábková et al., 2022; Schiffer et al., 2018). Including these phyla would have resulted in a searchable tree space with more than two million topologies, precluding exhaustive topology comparisons. The inclusion of even a single additional phylum would have increased our testable topologies to 945 rooted trees.

We interpret our results as showing that the five well-defined spiralian phyla (Annelida, Mollusca, Platyhelminthes, Nemertea and Brachiozoa) emerged in quick succession in a rapid or explosive radiation (Rokas and Carroll, 2006). Stronger signals within the wider Protostomia separate Ecdysozoa from Lophotrochozoa, and Gnathifera from the Spiralia. Within Spiralia evidence is mounting for lophophorate monophyly (Lewin et al., 2025; Lewin et al., 2026). While the expansion of completely sequenced genomes promises better data to address this problem, the short internodes we have observed predict that other genomic synapomorphies (e.g., unique macro and micro synteny patterns or informative examples of clade-specific gene presence/absence) are unlikely to have appeared in the short time frames we have described.

While beyond the scope of this work, the greater availability of genomes might help quantify the relative contribution of ILS and introgression to the high level of interphylum discordance identified in our tested datasets. Given that our inferred quartet scores for interphylum spiralian relationships are congruent with hard-polytomies (approximately 1:1:1 ratio, Fig. S1), and thus near to zero phylogenetic signal, we suspect the tested datasets (MARL and SERR) would not be sufficient to identify convincingly the predominant source of gene-tree discordance. It should be noted that spiralians are considerably older (>543.0 million years according to Carlisle et al. 2024) than the oldest ancient introgression events between metazoans identified to date (e.g., >200 million years for Odonata (Suvorov et al., 2022)).

The short internal branches and lack of signal suggest a short time between cladogenesis events and precious little opportunity for the evolution of major new characteristics, body plans or life cycles. Under this interpretation, it seems unlikely that characteristics shared by various combinations of spiralian phyla (shells, segmentation, larval types, chaetae) are synapomorphies of any superphyletic clades. A more plausible scenario involves some combination of loss in certain phyla of characters that had been present in a spiralian common ancestor and convergent or parallel evolution of characters in others. The lack of clarity as regards the evolutionary history of the characters possibly present in the stem-lineages of the Spiralia and the spiralian phyla make reconstruction of the evolution of these groups, as well as the interpretation of potential spiralian fossils, a difficult problem. The various Cambrian fossils showing mixed sets of characters (e.g. *Wiwaxia corrugata*, which may have a molluscan radula and annelidan chaetae (Zhang et al., 2015); *Pelagiella exigua*, with annelidan chaetae and a molluscan coiled shell (Landing et al., 2021); the tomotiid *Wufengella bengtsoni*, with chaetae, segmentation and a complex brachiopod-like mineralised skeleton (Guo et al., 2022)) could be interpreted as being on the stem lineage of one of the modern phyla and diverging before the loss (in the crown group) of one or more of the characters that we suggest were present in the spiralian ancestor.

The difficulties we have found are not restricted to Spiralia. Previous work, using both empirical and simulated data, has shown that, not only is systematic error widespread in metazoan data (Kapli et al., 2021; Kapli and Telford, 2020; Serra Silva et al., 2025; Song et al., 2023) but that even under conditions that mitigate these errors, it is often still not possible to distinguish between alternative topologies (Serra Silva et al., 2025). Our results add to a growing body of work suggesting that rapid cladogenesis might have been a common aspect of early metazoan evolution.

| Key Resources Table |  |  |  |  |
| --- | --- | --- | --- | --- |
| Reagent or resource | Designation | Source | Identifier | Additional information |
| Other | Alignments, phylogenetic analyses, simulations and ancillary scripts | DRYAD | https://doi.org/10.5061/dryad.280gb5n3j |  |
| Software and algorithms | IQ-TREE2 (v.2.3.5) | Minh et al., 2020 | http://www.iqtree.org/ |  |
| Software and algorithms | EDCluster | Schrempf et al., 2020 | https://github.com/dschrempf/EDCluster |  |
| Software and algorithms | ASTRAL-III (v.5.1.3) | Zhang et al., 2017 | https://github.com/smirarab/ASTRAL |  |
| Software and algorithms | MAFFT (v.7.455) | Katoh and Standley, 2013 | https://mafft.cbrc.jp/alignment/software/ |  |
| Software and algorithms | trimAl (v.1.4.rev15) | Capella-Gutiérrez et al., 2009 | https://vicfero.github.io/trimAl/ |  |
| Software and algorithms | Amino acid diversity tests | Giacomelli et al., 2022; Giacomelli et al., 2024 | https://github.com/mgiacom/tardigrades_catpmsf/tree/main |  |
| Software and algorithms | Additional data processing and analysis scripts | DRYAD | https://doi.org/10.5061/dryad.280gb5n3j |  |

## Materials and Methods

### Data collection

All analyses were run on two datasets derived from independent studies on animal relationships, Marlétaz et al. (2019) and Serra Silva et al. (2025)—referred throughout as MARL and SERR, respectively. To limit computational burden, the taxon sampling of both datasets was reduced by removal of the majority of non-lophotrochozoan taxa and spiralian phyla outside of the five focal groups (Annelida, Brachiozoa, Mollusca, Nemertea and Platyhelminthes). Highly labile spiralian and gnathiferan taxa were also removed prior to taxon sampling reduction, based on extensive paraphyly with non-target phyla in filtering supertree analyses (e.g. *Owenia fusiformis* in SERR).

After taxonomic filtering, we retained loci with a minimum of 75% taxon coverage and realigned them with mafft v.7.455 (Katoh and Standley, 2013) under automatic algorithm selection. The resulting alignments were run through trimAl v.1.4.rev15 (Capella-Gutiérrez et al., 2009) to remove gap-rich regions (‘-gappyout’). To further limit computational burden, we reduced the MARL matrix from c.335,000 amino acid positions to approximately the same length and proportion of missing data as the SERR matrix by retaining loci with at least 89% taxon-coverage. Results are based on two concatenated matrices with 82 taxa, 66,760 amino acid positions and 69% coverage for SERR and 56 taxa, 70,629 amino acid positions and 65% coverage for MARL.

### Topology-scoring on fixed-taxon sampling matrices

For each fixed-taxon dataset, we computed log-likelihood scores for each of the possible 105 rooted Spiralia trees using IQ-Tree v.2.3.5 (Minh et al., 2020), under the site-homogeneous LG+G4+F and site-heterogeneous EDM+G4+F models. We selected the ‘-wslr’ flag to output site-specific log-likelihoods to a file. The dataset-specific 128 category EDM matrices were obtained following the protocol in Schrempf et al. (2020). Intra-phylum relationships were fixed for all topology-scoring analyses, based on unconstrained tree searches of the filtered matrices under LG+F+G.

Following the resampling estimated log-likelihood (RELL) method (Kishino et al., 1990), we used the IQ-Tree-computed site-specific log-likelihoods to generate 10,000 non-parametric bootstrap replicates per topology. We repeated the resampling for each dataset and model.

We repeated these analyses on fixed-taxon matrices with all outgroups removed (Cnidaria, Chordata, Ambulacraria, Ecdysozoa and Gnathifera) and topologies restricted to the 15 unrooted topologies depicted in Fig. 1.

### Branch-length calculation

From the fixed-taxon sampling log-likelihood scoring analyses on the 105 rooted topologies, we recorded the length of each of the three internal branches connecting the five Spiralia clades (Annelida, Brachiozoa, Mollusca, Nemertea and Platyhelminthes), as well as those branches leading to each phylum and to Gnathifera, Deuterostomia and Protostomia (script in DRYAD repository doi.org/10.5061/dryad.280gb5n3j).

### Randomised taxon-jackknifing

We generated 100 taxon-jackknife replicates for each fixed sampling matrix (with or without outgroups). For the rooted topology analyses, each replicate included three randomly drawn taxa for each Spiralia phylum (Annelida, Brachiopoda+Phoronida, Mollusca, Nemertea and Platyhelminthes) and Gnathifera, with the non-Lophotrochozoa taxa remaining unchanged for all replicates. This generated 20 matrices with 31 taxa for SERR, and 30 taxa for MARL. All replicates were run through IQ-Tree under the site-homogeneous LG+G4+F and the site-heterogeneous EDM+G4+F models and each of the 105 fixed rooted topologies.

For the ingroup-only analyses, we randomly sampled three representatives of each spiralian phylum, generating 100 matrices with 15 taxa for both empirical datasets. All replicates were run through IQ-Tree under the site-homogeneous LG+G4+F and the site-heterogeneous EDM+G4+F models and each of the 15 unrooted topologies.

### Simulations under an unresolved Spiralia tree

To test whether any of the observed support for rooting Spiralia on the platyhelminth branch and for unrooted uT4 in the ingroup-only analyses might be due to unaccounted for compositional heterogeneity across sites, we simulated data on an unrooted Spiralia tree with all inter-phylum branches collapsed (star-tree) and on a rooted Lophotrochozoa star-tree (resolved outgroups and Gnathifera subtree with internally unresolved Spiralia). For each dataset, we used IQ-Tree’s inbuilt AliSim (Ly-Trong et al., 2022), which allows for polytomous trees, to generate 100 simulated alignments under the site-heterogeneous EDM+G4. The parameter ‘+F’ was omitted as it was hard-coded in AliSim for the IQ-Tree version used.

All simulated alignments were scored under the LG+G4 model (Lophotrochozoa and outgroups—105 rooted trees; Spiralia-only—15 unrooted trees) and, to limit computational burden, a selection of 10 alignments generated under the site-heterogeneous EDM model were scored under the ‘correct’ model.

### Statistical tests on topology ranking

For all RELL, randomised taxon-jackknives and likelihood scoring on simulated trees, we summarised the per topology log-likelihoods with boxplots and tested for significant differences in per topology support with a series of multi-sample non-parametric Kruskal-Wallis H-tests. For samples with significant (α = 0.05) differences, *post hoc* Welch’s two-sample t-tests were performed to identify significantly different pairs of topologies.

### Discordance-aware analyses

To test for gene-tree discordance, we inferred ML gene-trees for each individual alignment from both the SERR and MARL datasets under the LG+G4+F model. The resulting gene-trees were used as input for an Astral-III v.5.1.3 (Zhang et al., 2017) summary tree. We used the ‘-t 32’ option to obtain the quartet scores (QS) of alternative quartet topologies. See supplementary text and figure S1 for results.

### Compositional diversity of posterior samples

We used the simulation-based z-score amino acid diversity tests described in Giacomelli et al. (2022; 2024) to compare the model-fit of 128-EDM+G4 and LG+G4 to both the SERR and MARL datasets. We simulated data with IQ-Tree’s AliSim, with the same parameters described above but using rooted T22 as the guide tree. Results provided in table S2.

## Acknowledgments

We thank Paschalia Kapli (NHM) for comments. This work was funded by Leverhulme Trust Research Grant RPG-2021-433 (ASS, MJT).

## Author contributions

Conceptualization: ASS, MJT; Methodology: ASS, MJT; Investigation: ASS; Data curation: ASS; Visualization: ASS, MJT; Funding acquisition: MJT; Writing – original draft: ASS, MJT; Writing – review and editing: ASS, MJT

## Competing interests

Authors declare that they have no competing interests.

## Data and materials availability

All data are available in the main text, supplementary materials or the DRYAD repository doi.org/10.5061/dryad.280gb5n3j.

## Supplementary Materials

Supplementary Text

Figs. S1 to S2

Tables S1 to S11

Data S1

---

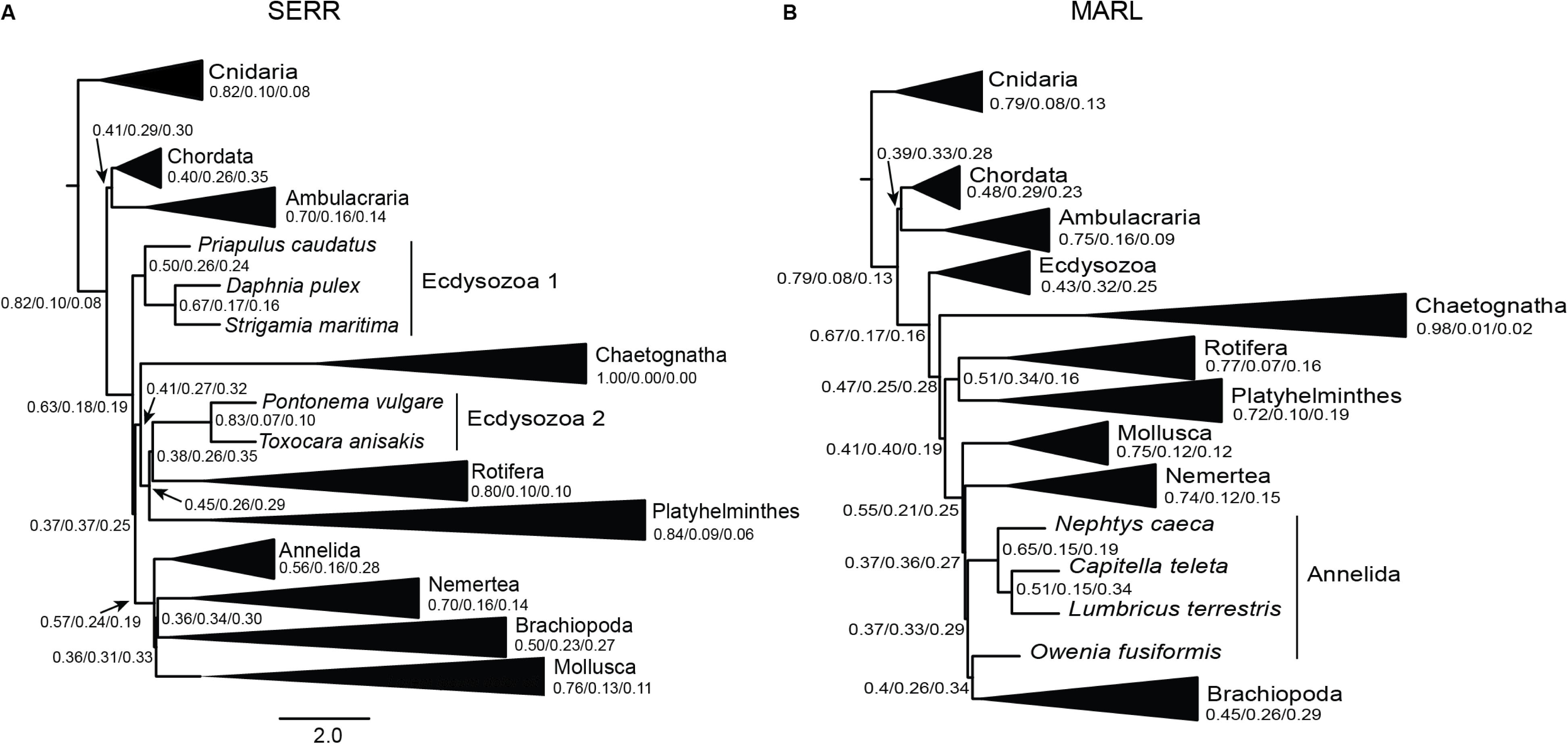

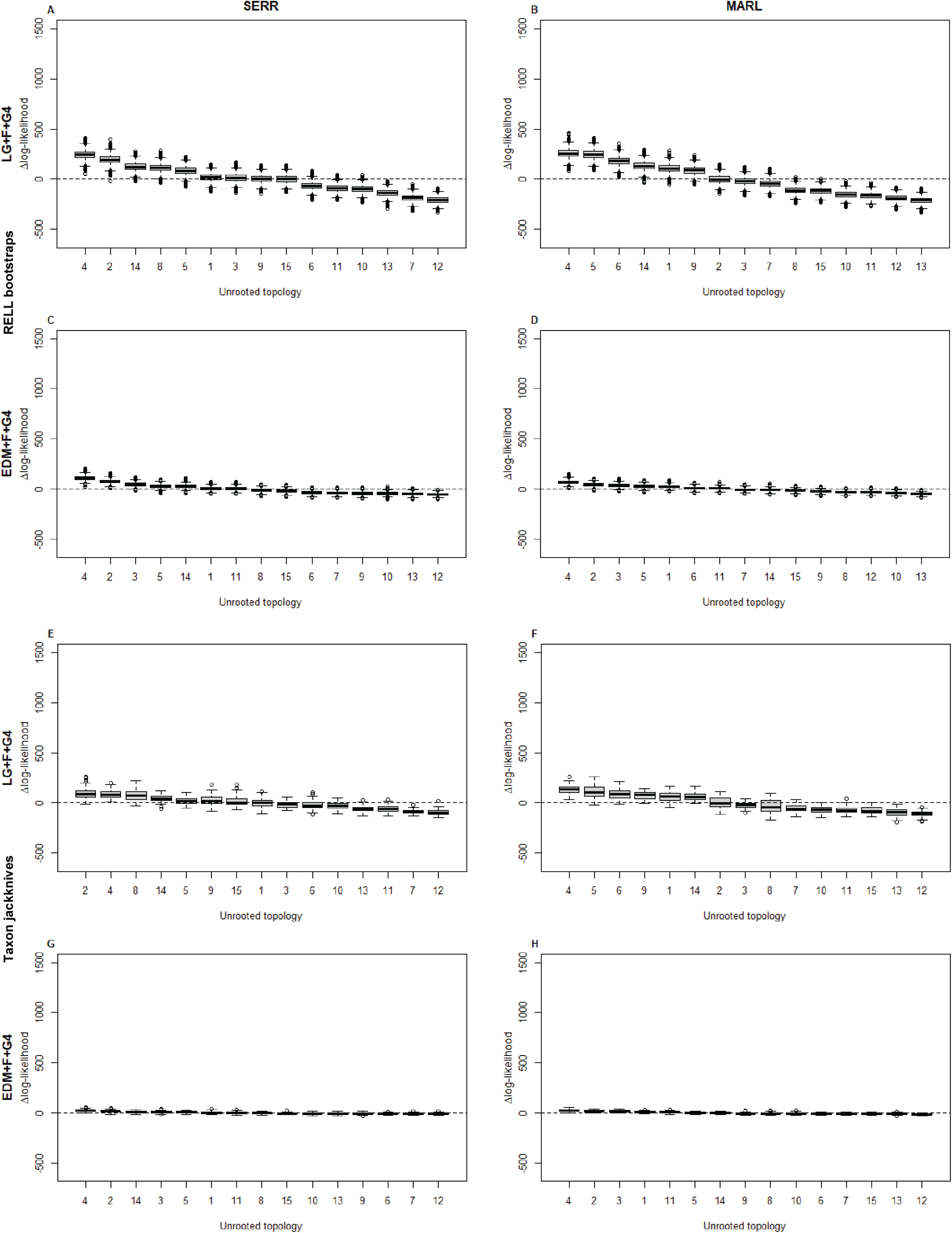

## Notes

### Competing Interest Statement

The authors have declared no competing interest.

### Summary of Updates

We have responded to the reviews organised by the journal elife and the changes can be found on the publishers website accompanying this article.

