## Supplementary info for "Are interphylum spiralian relationships resolvable?"

**The file includes:**

Supplementary text
Figs. S1 to S2
Tables S1 to S11

**Supplementary text**

Notes on nomenclature

In the main text of this paper, we used the name Lophotrochozoa (Halanych et al., 1995) to refer to the sister group of the Ecdysozoa. We consider Lophotrochozoa to include two clades: the first, Spiralia, includes the spirally cleaving Mollusca, Annelida, Nemertea, Platyhelminthes, Brachiopoda and Phoronida, as well as Entoprocta, Gastrotricha and Ectoprocta/Bryozoa; the second, Gnathifera, unites mostly tiny, jaw-bearing animals—Chaetognatha, Gnathostomulida (possibly also spirally cleaving according to Riedl (1969)), Rotifera and Micrognathozoa (Marlétaz et al., 2019; Vinther & Parry, 2019). Given the often-interchangeable use of the names Lophotrochozoa and Spiralia, we provide an abridged history of the uses of these names.

The name 'Spiralia' was first used by Schleip (1929) as a descriptive term for a subset of the apparently disparate groups with this pattern of cell division (specifically the only taxa Schleip discusses are polyclad flatworms, nemerteans, annelids and molluscs). 'Spiralia' was later adopted by some authors as a clade name that described various groupings of phyla—most often encompassing most or all of what we now refer to as Protostomia (the majority of which do not have spiral cleavage) and in all cases including the Arthropoda alongside Annelida (within the Articulata).

The name first given to the clade that is the sister group of the Ecdysozoa—the clade we now recognise as containing the spiral cleaving taxa but not, importantly, the arthropods and several other non-spirally cleaving protostome phyla—was 'Lophotrochozoa' (Halanych et al., 1995). This clade had never been proposed prior to being recognised using molecular data. The widely accepted form of Lophotrochozoa (barring then-unsampled Gnathostomulida and Micrognathozoa) was explicitly outlined as long ago as 1997 (Aguinaldo et al., 1997) as including: Chaetognatha, Platyhelminthes, Nemertea, Mollusca, Acanthocephala, Gastrotricha, Rotifera, Phoronida, Ectoprocta, Brachiopoda and Annelida (within which we now place several other lophotrochozoan 'phyla' namely Sipuncula, Pogonophora, Echiura and Vestimentifera). The name Lophotrochozoa had been in wide use and its application unambiguous until Giribet (Giribet, 2002, 2008; Giribet et al., 2009) equated the old name 'Spiralia' to the distinct, clearly defined and already named Lophotrochozoa. This has led to inevitable confusion and inconsistent use of the superphyletic names Lophotrochozoa and Spiralia.

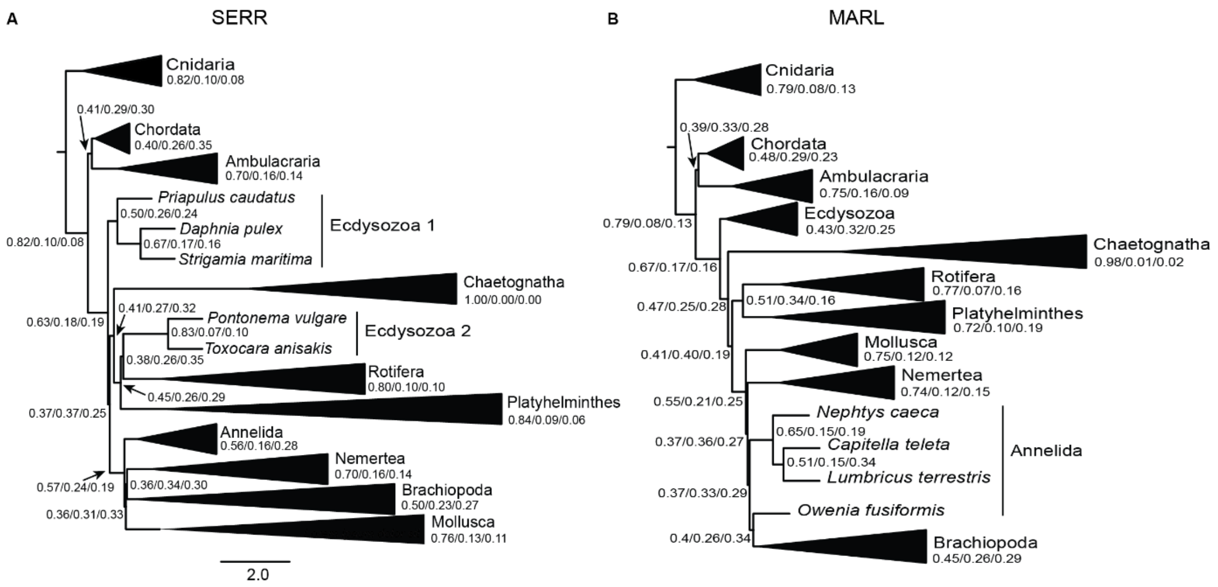


**Fig. S1.**

Discordance-aware summary tree analyses show extensive gene-tree conflict within Spiralia. For SERR (A), the Astral analysis recovers all spiralian phyla as monophyletic but no internal Spiralia branch with quartet scores (QS) over 0.50. The exception is the branch separating Platyhelminthes+outgroups from the remaining Spiralia phyla; however, Platyhelminthes are recovered within paraphyletic Gnathifera. For MARL, the Astral analysis yields very similar results, except that Annelida is not monophyletic. Branch-lengths are in coalescent units and monophyletic phyla have been collapsed, with the quartet scores (Q1/Q2/Q3) for the branch subtending each phylum listed underneath the phylum names. Detailed quartet scores can be found in the Astral.zip folder in the Dryad data repository (https://datadryad.org/downloads/file_stream/4710539).

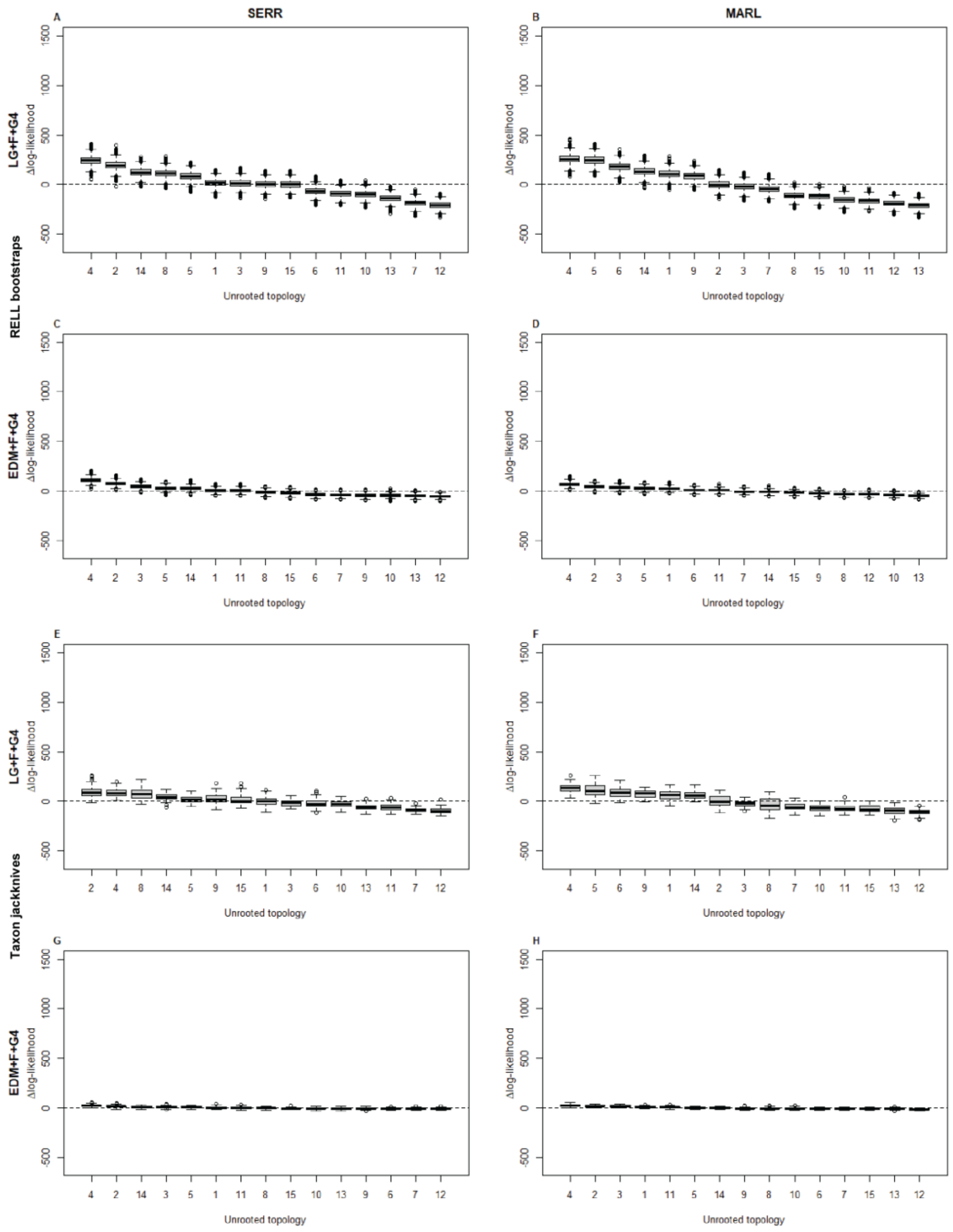


**Fig. S2.**

Relative median RELL pseudo-bootstraps (**A**–**D**) and randomised taxon-jackknives (**E**–**H**) log-likelihood ranks for all unrooted trees, see tables S8 and S9 for all ranks. Topologies ranked by their difference in log-likelihood to the mean log-likelihood of each RELL or jackknife distribution. The RELL analyses of both data sets under the site-homogeneous LG+F+G4 model (**A**, **B**) occupy the widest ranges of log-likelihoods, which is consistent with the statistically significant differences found by the Kruskal-Wallis test. The RELL analyses under the site-heterogeneous EDM+F+G4 model (**C**, **D**) occupy a much narrower range of log-likelihoods. The jackknife analyses show a similar trend, wider range of median log-likelihood scores under LG+F+G4 (**E**, **F**) than under EDM+F+G4 (**G**, **H**); however, the log-likelihood ranges of the jackknives under LG+F+G4 are very similar to that occupied by RELL under EDM+F+G4, consistent with the lack of statistically significant differences between topologies in the jackknife analyses.

**Table S1.**

Median branch lengths for the 105 rooted topologies. Phylum-level branch lengths shown in Fig. 2.

| Branch | SERR+LG | SERR+EDM | MARL+LG | MARL+EDM |
|---|---|---|---|---|
| Deuterostomia | 0.03139 | 0.02812 | 0.02305 | 0.01782 |
| Ecdysozoa | 0.04762 | 0.06257 | 0.03835 | 0.03941 |
| Lophotrochozoa | 0.02440 | 0.02607 | 0.02382 | 0.02613 |
| Protostomia | 0.06420 | 0.09325 | 0.04852 | 0.06641 |
| Internal | 0.01616 | 0.01393 | 0.01316 | 0.009462 |

**Table S2.**

Fixed-taxon RELL pseudo-bootstraps rankings for 105 rooted topologies.

| Topology | SERR+LG | SERR+EDM | MARL+LG | MARL+EDM |
|---|---|---|---|---|
| 1 | 40 | 34 | 26 | 12 |
| 2 | 5 | 3 | 2 | 8 |
| 3 | 38 | 35 | 15 | 7 |
| 4 | 73 | 77 | 46 | 27 |
| 5 | 71 | 57 | 38 | 21 |
| 6 | 92 | 81 | 77 | 77 |
| 7 | 101 | 91 | 61 | 80 |
| 8 | 56 | 66 | 80 | 24 |
| 9 | 35 | 31 | 48 | 6 |
| 10 | 27 | 22 | 40 | 11 |
| 11 | 41 | 42 | 56 | 62 |
| 12 | 19 | 21 | 51 | 55 |
| 13 | 30 | 24 | 66 | 57 |
| 14 | 2 | 2 | 13 | 40 |
| 15 | 72 | 70 | 68 | 23 |
| 16 | 50 | 36 | 39 | 5 |
| 17 | 33 | 23 | 34 | 9 |
| 18 | 4 | 1 | 6 | 10 |
| 19 | 43 | 25 | 42 | 19 |
| 20 | 77 | 54 | 70 | 71 |
| 21 | 64 | 45 | 82 | 67 |
| 22 | 39 | 38 | 37 | 4 |
| 23 | 22 | 18 | 22 | 1 |
| 24 | 17 | 12 | 20 | 2 |
| 25 | 21 | 16 | 32 | 18 |
| 26 | 18 | 13 | 12 | 15 |
| 27 | 45 | 26 | 35 | 49 |
| 28 | 6 | 4 | 5 | 29 |
| 29 | 57 | 62 | 53 | 33 |
| 30 | 46 | 47 | 52 | 37 |
| 31 | 34 | 28 | 25 | 14 |
| 32 | 23 | 19 | 14 | 3 |
| 33 | 20 | 14 | 8 | 13 |
| 34 | 55 | 33 | 29 | 51 |
| 35 | 10 | 5 | 3 | 30 |
| 36 | 97 | 96 | 69 | 59 |
| 37 | 90 | 86 | 71 | 61 |
| 38 | 75 | 60 | 33 | 31 |
| 39 | 78 | 68 | 31 | 52 |
| 40 | 48 | 40 | 10 | 20 |
| 41 | 49 | 39 | 21 | 26 |
| 42 | 7 | 6 | 1 | 22 |

| Topology | SERR+LG | SERR+EDM | MARL+LG | MARL+EDM |
|---|---|---|---|---|
| 43 | 100 | 92 | 95 | 72 |
| 44 | 89 | 78 | 101 | 79 |
| 45 | 70 | 53 | 49 | 43 |
| 46 | 13 | 8 | 9 | 28 |
| 47 | 74 | 51 | 59 | 53 |
| 48 | 93 | 69 | 65 | 68 |
| 49 | 67 | 46 | 41 | 32 |
| 50 | 82 | 102 | 99 | 99 |
| 51 | 81 | 100 | 85 | 103 |
| 52 | 47 | 88 | 67 | 87 |
| 53 | 29 | 49 | 45 | 44 |
| 54 | 16 | 32 | 58 | 83 |
| 55 | 26 | 43 | 72 | 88 |
| 56 | 1 | 7 | 19 | 66 |
| 57 | 102 | 104 | 73 | 86 |
| 58 | 103 | 105 | 60 | 94 |
| 59 | 66 | 95 | 44 | 69 |
| 60 | 52 | 75 | 50 | 70 |
| 61 | 36 | 55 | 16 | 34 |
| 62 | 37 | 52 | 28 | 42 |
| 63 | 3 | 9 | 4 | 35 |
| 64 | 98 | 103 | 96 | 101 |
| 65 | 96 | 101 | 86 | 104 |
| 66 | 60 | 90 | 64 | 90 |
| 67 | 8 | 20 | 11 | 65 |
| 68 | 54 | 89 | 63 | 84 |
| 69 | 51 | 63 | 55 | 46 |
| 70 | 59 | 76 | 90 | 91 |
| 71 | 65 | 64 | 74 | 41 |
| 72 | 11 | 11 | 17 | 25 |
| 73 | 63 | 61 | 79 | 45 |
| 74 | 58 | 44 | 54 | 17 |
| 75 | 84 | 59 | 75 | 38 |
| 76 | 99 | 82 | 105 | 98 |
| 77 | 105 | 93 | 102 | 102 |
| 78 | 88 | 83 | 94 | 74 |
| 79 | 14 | 17 | 27 | 58 |
| 80 | 85 | 79 | 100 | 82 |
| 81 | 91 | 80 | 103 | 93 |
| 82 | 94 | 71 | 97 | 76 |
| 83 | 86 | 65 | 78 | 56 |
| 84 | 104 | 87 | 98 | 96 |
| 85 | 87 | 98 | 89 | 95 |
| 86 | 15 | 27 | 24 | 78 |
| 87 | 83 | 97 | 92 | 100 |

| Topology | SERR+LG | SERR+EDM | MARL+LG | MARL+EDM |
|---|---|---|---|---|
| 88 | 95 | 99 | 88 | 105 |
| 89 | 80 | 94 | 87 | 92 |
| 90 | 69 | 72 | 81 | 54 |
| 91 | 79 | 84 | 104 | 97 |
| 92 | 25 | 29 | 30 | 16 |
| 93 | 31 | 41 | 57 | 47 |
| 94 | 32 | 56 | 36 | 39 |
| 95 | 42 | 74 | 43 | 48 |
| 96 | 24 | 30 | 18 | 36 |
| 97 | 62 | 67 | 47 | 85 |
| 98 | 12 | 15 | 7 | 63 |
| 99 | 53 | 50 | 62 | 50 |
| 100 | 76 | 73 | 84 | 89 |
| 101 | 61 | 58 | 83 | 64 |
| 102 | 68 | 85 | 93 | 75 |
| 103 | 28 | 37 | 76 | 73 |
| 104 | 44 | 48 | 91 | 81 |
| 105 | 9 | 10 | 23 | 60 |

**Table S3.**

Fixed-taxon RELL pseudo-bootstraps best-ranked topology frequencies for 105 rooted topologies. Omitted topologies were *never* recovered as the best-ranked tree.

| Topology | SERR+LG | SERR+EDM | MARL+LG | MARL+EDM |
|---|---|---|---|---|
| 2 | 8.4 | 11.96 | 27.62 | 0.43 |
| 3 | 0 | 0 | 0 | 0.59 |
| 14 | 8.09 | 15.74 | 0 | 0 |
| 18 | 11.23 | 51.78 | 1.72 | 0.34 |
| 22 | 0 | 0 | 0 | 0.04 |
| 23 | 0 | 0 | 0 | 57.2 |
| 24 | 0 | 0.28 | 0 | 36.52 |
| 26 | 0 | 0.1 | 0.01 | 0.05 |
| 28 | 0.41 | 3.8 | 4.72 | 0 |
| 29 | 0 | 0 | 0 | 0.01 |
| 31 | 0 | 0 | 0 | 0.67 |
| 32 | 0 | 0 | 0 | 3.53 |
| 33 | 0 | 0.04 | 0.36 | 0.5 |
| 35 | 0.26 | 6.15 | 13.15 | 0 |
| 38 | 0 | 0 | 0 | 0.01 |
| 39 | 0 | 0 | 0 | 0 |
| 40 | 0 | 0 | 0 | 0.01 |
| 42 | 0.34 | 5.26 | 50.33 | 0.05 |
| 46 | 0.01 | 1.49 | 0 | 0.02 |
| 56 | 63.53 | 2.23 | 0 | 0 |
| 61 | 0 | 0 | 0 | 0.02 |
| 63 | 7.65 | 0.54 | 2.03 | 0 |
| 67 | 0.03 | 0 | 0 | 0 |
| 72 | 0 | 0.07 | 0 | 0 |
| 92 | 0 | 0 | 0 | 0.01 |
| 98 | 0 | 0 | 0.06 | 0 |
| 105 | 0.05 | 0.5 | 0 | 0 |

**Table S4.**

Randomised taxon-jackknife rankings for 105 rooted topologies.

| Topology | SERR+LG | SERR+EDM | MARL+LG | MARL+EDM |
|---|---|---|---|---|
| 1 | 39 | 18 | 36 | 19 |
| 2 | 7 | 15 | 3 | 24 |
| 3 | 22 | 14 | 6 | 10 |
| 4 | 63 | 64 | 30 | 13 |
| 5 | 54 | 13 | 28 | 17 |
| 6 | 80 | 54 | 75 | 83 |
| 7 | 100 | 88 | 65 | 72 |
| 8 | 58 | 45 | 85 | 11 |
| 9 | 33 | 12 | 52 | 6 |
| 10 | 20 | 2 | 35 | 4 |
| 11 | 48 | 41 | 50 | 53 |
| 12 | 15 | 7 | 45 | 50 |
| 13 | 34 | 16 | 77 | 69 |
| 14 | 4 | 9 | 19 | 70 |
| 15 | 77 | 58 | 71 | 12 |
| 16 | 52 | 11 | 48 | 7 |
| 17 | 24 | 4 | 26 | 5 |
| 18 | 2 | 3 | 1 | 14 |
| 19 | 35 | 5 | 43 | 20 |
| 20 | 91 | 61 | 79 | 76 |
| 21 | 67 | 27 | 91 | 77 |
| 22 | 55 | 33 | 38 | 3 |
| 23 | 38 | 6 | 22 | 2 |
| 24 | 19 | 1 | 11 | 1 |
| 25 | 27 | 8 | 31 | 23 |
| 26 | 23 | 10 | 9 | 16 |
| 27 | 56 | 37 | 47 | 60 |
| 28 | 9 | 24 | 12 | 54 |
| 29 | 72 | 59 | 63 | 26 |
| 30 | 61 | 40 | 64 | 40 |
| 31 | 43 | 22 | 20 | 9 |
| 32 | 31 | 31 | 15 | 8 |
| 33 | 36 | 65 | 10 | 18 |
| 34 | 75 | 87 | 44 | 67 |
| 35 | 13 | 80 | 13 | 58 |
| 36 | 93 | 81 | 69 | 46 |
| 37 | 79 | 63 | 78 | 63 |
| 38 | 64 | 55 | 25 | 25 |
| 39 | 92 | 92 | 27 | 42 |
| 40 | 30 | 56 | 7 | 22 |
| 41 | 42 | 46 | 32 | 28 |
| 42 | 8 | 42 | 4 | 37 |

| Topology | SERR+LG | SERR+EDM | MARL+LG | MARL+EDM |
|---|---|---|---|---|
| 43 | 96 | 76 | 96 | 71 |
| 44 | 90 | 70 | 102 | 90 |
| 45 | 60 | 38 | 58 | 44 |
| 46 | 11 | 26 | 16 | 52 |
| 47 | 68 | 47 | 67 | 57 |
| 48 | 101 | 93 | 72 | 78 |
| 49 | 69 | 53 | 57 | 30 |
| 50 | 82 | 100 | 99 | 85 |
| 51 | 83 | 104 | 86 | 94 |
| 52 | 29 | 69 | 54 | 56 |
| 53 | 21 | 21 | 39 | 32 |
| 54 | 12 | 17 | 49 | 66 |
| 55 | 28 | 30 | 76 | 82 |
| 56 | 1 | 20 | 17 | 84 |
| 57 | 95 | 101 | 60 | 61 |
| 58 | 99 | 105 | 55 | 75 |
| 59 | 46 | 77 | 23 | 38 |
| 60 | 37 | 50 | 42 | 55 |
| 61 | 16 | 39 | 5 | 33 |
| 62 | 26 | 34 | 24 | 43 |
| 63 | 3 | 32 | 2 | 50 |
| 64 | 97 | 98 | 92 | 87 |
| 65 | 102 | 102 | 88 | 95 |
| 66 | 44 | 71 | 46 | 68 |
| 67 | 6 | 60 | 8 | 65 |
| 68 | 45 | 68 | 51 | 64 |
| 69 | 50 | 35 | 56 | 39 |
| 70 | 49 | 43 | 90 | 89 |
| 71 | 70 | 75 | 70 | 34 |
| 72 | 10 | 57 | 21 | 41 |
| 73 | 66 | 79 | 61 | 45 |
| 74 | 53 | 25 | 53 | 21 |
| 75 | 71 | 23 | 62 | 36 |
| 76 | 98 | 62 | 104 | 105 |
| 77 | 105 | 89 | 100 | 102 |
| 78 | 87 | 84 | 98 | 74 |
| 79 | 14 | 78 | 41 | 91 |
| 80 | 81 | 86 | 93 | 93 |
| 81 | 84 | 74 | 103 | 104 |
| 82 | 88 | 66 | 95 | 79 |
| 83 | 86 | 72 | 80 | 49 |
| 84 | 104 | 95 | 97 | 92 |
| 85 | 89 | 97 | 87 | 96 |
| 86 | 17 | 94 | 34 | 100 |
| 87 | 85 | 99 | 82 | 101 |

| Topology | SERR+LG | SERR+EDM | MARL+LG | MARL+EDM |
|---|---|---|---|---|
| 88 | 103 | 103 | 84 | 103 |
| 89 | 73 | 83 | 73 | 81 |
| 90 | 76 | 44 | 81 | 48 |
| 91 | 78 | 52 | 105 | 99 |
| 92 | 40 | 19 | 33 | 15 |
| 93 | 47 | 28 | 66 | 47 |
| 94 | 41 | 49 | 29 | 29 |
| 95 | 62 | 85 | 40 | 31 |
| 96 | 32 | 73 | 14 | 27 |
| 97 | 74 | 96 | 59 | 88 |
| 98 | 18 | 90 | 18 | 86 |
| 99 | 57 | 67 | 68 | 35 |
| 100 | 94 | 91 | 89 | 73 |
| 101 | 59 | 51 | 83 | 59 |
| 102 | 65 | 82 | 94 | 62 |
| 103 | 25 | 29 | 74 | 80 |
| 104 | 51 | 48 | 101 | 98 |
| 105 | 5 | 36 | 37 | 97 |

**Table S5.**

Randomised taxon-jackknife best-ranked topology frequencies for 105 rooted topologies. Omitted topologies were *never* recovered as the best-ranked tree.

| Topology | SERR+LG | SERR+EDM | MARL+LG | MARL+EDM |
|---|---|---|---|---|
| 2 | 15 | 0 | 0 | 0 |
| 8 | 0 | 0 | 0 | 5 |
| 9 | 0 | 0 | 0 | 0 |
| 10 | 0 | 10 | 0 | 5 |
| 12 | 0 | 5 | 0 | 0 |
| 14 | 5 | 5 | 0 | 0 |
| 17 | 0 | 5 | 0 | 0 |
| 18 | 30 | 5 | 25 | 0 |
| 23 | 0 | 5 | 0 | 30 |
| 24 | 0 | 40 | 0 | 55 |
| 26 | 0 | 5 | 5 | 0 |
| 31 | 0 | 5 | 0 | 0 |
| 32 | 0 | 5 | 0 | 5 |
| 33 | 0 | 0 | 15 | 0 |
| 40 | 0 | 0 | 10 | 0 |
| 42 | 0 | 0 | 5 | 0 |
| 54 | 10 | 0 | 0 | 0 |
| 56 | 30 | 0 | 0 | 0 |
| 61 | 0 | 0 | 10 | 0 |
| 63 | 10 | 0 | 20 | 0 |
| 67 | 0 | 0 | 10 | 0 |
| 92 | 0 | 5 | 0 | 0 |
| 103 | 0 | 5 | 0 | 0 |

**Table S6.**

Simulations on rooted Spiralia star-tree rankings for 105 rooted topologies. For columns SERR+EDM and MARL+EDM, proportions are based on 10 alignments.

| Topology | SERR+LG EDM sim | SERR+EDM | MARL+LG EDM sim | MARL+EDM |
|---|---|---|---|---|
| 1 | 24 | 9 | 24 | 71 |
| 2 | 4 | 2 | 4 | 7 |
| 3 | 21 | 41 | 18 | 34 |
| 4 | 57 | 44 | 37 | 44 |
| 5 | 54 | 89 | 36 | 46 |
| 6 | 69 | 103 | 70 | 47 |
| 7 | 81 | 91 | 56 | 61 |
| 8 | 95 | 86 | 105 | 93 |
| 9 | 64 | 90 | 101 | 78 |
| 10 | 49 | 46 | 93 | 80 |
| 11 | 51 | 61 | 86 | 87 |
| 12 | 23 | 19 | 58 | 56 |
| 13 | 26 | 53 | 73 | 86 |
| 14 | 6 | 5 | 14 | 8 |
| 15 | 94 | 99 | 81 | 99 |
| 16 | 68 | 57 | 77 | 92 |
| 17 | 47 | 95 | 55 | 102 |
| 18 | 10 | 31 | 9 | 11 |
| 19 | 48 | 97 | 48 | 100 |
| 20 | 73 | 77 | 88 | 98 |
| 21 | 62 | 73 | 95 | 96 |
| 22 | 103 | 81 | 74 | 21 |
| 23 | 78 | 87 | 68 | 28 |
| 24 | 55 | 8 | 49 | 24 |
| 25 | 53 | 22 | 50 | 9 |
| 26 | 37 | 3 | 21 | 32 |
| 27 | 39 | 42 | 33 | 101 |
| 28 | 11 | 47 | 6 | 16 |
| 29 | 102 | 29 | 65 | 23 |
| 30 | 90 | 58 | 66 | 20 |
| 31 | 50 | 71 | 29 | 25 |
| 32 | 46 | 70 | 32 | 35 |
| 33 | 34 | 50 | 17 | 48 |
| 34 | 38 | 79 | 23 | 105 |
| 35 | 8 | 48 | 2 | 19 |
| 36 | 87 | 65 | 57 | 42 |
| 37 | 67 | 26 | 62 | 27 |
| 38 | 41 | 83 | 27 | 50 |
| 39 | 43 | 76 | 30 | 55 |
| 40 | 19 | 68 | 16 | 37 |

| Topology | SERR+LG EDM sim | SERR+EDM | MARL+LG EDM sim | MARL+EDM |
|---|---|---|---|---|
| 41 | 22 | 40 | 22 | 82 |
| 42 | 3 | 7 | 1 | 1 |
| 43 | 96 | 62 | 98 | 75 |
| 44 | 79 | 45 | 99 | 60 |
| 45 | 42 | 78 | 39 | 79 |
| 46 | 9 | 60 | 7 | 13 |
| 47 | 44 | 98 | 45 | 90 |
| 48 | 61 | 94 | 63 | 103 |
| 49 | 52 | 80 | 61 | 76 |
| 50 | 91 | 34 | 104 | 64 |
| 51 | 85 | 67 | 103 | 72 |
| 52 | 31 | 20 | 84 | 67 |
| 53 | 30 | 35 | 78 | 77 |
| 54 | 17 | 10 | 53 | 69 |
| 55 | 20 | 39 | 67 | 65 |
| 56 | 2 | 4 | 13 | 12 |
| 57 | 80 | 36 | 71 | 51 |
| 58 | 74 | 75 | 64 | 59 |
| 59 | 29 | 16 | 43 | 54 |
| 60 | 28 | 24 | 44 | 41 |
| 61 | 16 | 37 | 19 | 45 |
| 62 | 18 | 13 | 26 | 81 |
| 63 | 1 | 1 | 3 | 2 |
| 64 | 99 | 28 | 87 | 95 |
| 65 | 98 | 84 | 85 | 97 |
| 66 | 32 | 12 | 54 | 83 |
| 67 | 5 | 11 | 10 | 18 |
| 68 | 33 | 82 | 47 | 94 |
| 69 | 35 | 43 | 83 | 88 |
| 70 | 36 | 30 | 92 | 85 |
| 71 | 60 | 100 | 35 | 84 |
| 72 | 13 | 105 | 8 | 3 |
| 73 | 56 | 21 | 25 | 43 |
| 74 | 66 | 14 | 51 | 53 |
| 75 | 82 | 23 | 46 | 49 |
| 76 | 100 | 69 | 91 | 29 |
| 77 | 105 | 74 | 82 | 70 |
| 78 | 63 | 88 | 42 | 74 |
| 79 | 14 | 63 | 12 | 6 |
| 80 | 59 | 59 | 34 | 36 |
| 81 | 76 | 25 | 80 | 15 |
| 82 | 84 | 64 | 76 | 39 |
| 83 | 92 | 52 | 90 | 58 |
| 84 | 104 | 85 | 94 | 68 |

| Topology | SERR+LG EDM sim | SERR+EDM | MARL+LG EDM sim | MARL+EDM |
|---|---|---|---|---|
| 85 | 72 | 96 | 38 | 89 |
| 86 | 15 | 101 | 11 | 5 |
| 87 | 70 | 56 | 28 | 40 |
| 88 | 101 | 51 | 59 | 63 |
| 89 | 89 | 15 | 52 | 38 |
| 90 | 86 | 38 | 79 | 33 |
| 91 | 93 | 33 | 89 | 30 |
| 92 | 75 | 72 | 60 | 14 |
| 93 | 83 | 17 | 72 | 10 |
| 94 | 77 | 32 | 40 | 17 |
| 95 | 97 | 49 | 41 | 26 |
| 96 | 40 | 18 | 20 | 52 |
| 97 | 45 | 93 | 31 | 104 |
| 98 | 12 | 66 | 5 | 22 |
| 99 | 65 | 92 | 100 | 73 |
| 100 | 88 | 102 | 102 | 91 |
| 101 | 58 | 104 | 96 | 66 |
| 102 | 71 | 55 | 97 | 57 |
| 103 | 25 | 27 | 69 | 31 |
| 104 | 27 | 54 | 75 | 62 |
| 105 | 7 | 6 | 15 | 4 |

**Table S7.**

Simulations on rooted Spiralia star-tree best-ranked topology frequencies for 105 rooted topologies. For columns SERR+EDM and MARL+EDM, proportions are based on 10 alignments. Omitted topologies were *never* recovered as the best-ranked tree.

| Topology | SERR+LG EDM sim | SERR+EDM | MARL+LG EDM sim | MARL+EDM |
|---|---|---|---|---|
| 10 | 0 | 0 | 0 | 20 |
| 18 | 0 | 20 | 0 | 0 |
| 23 | 0 | 20 | 0 | 0 |
| 28 | 0 | 0 | 15 | 0 |
| 33 | 0 | 0 | 1 | 0 |
| 35 | 0 | 0 | 84 | 0 |
| 49 | 2 | 0 | 0 | 0 |
| 56 | 98 | 0 | 0 | 0 |
| 60 | 0 | 20 | 0 | 0 |
| 72 | 0 | 0 | 0 | 20 |
| 85 | 0 | 0 | 0 | 20 |
| 86 | 0 | 0 | 0 | 20 |
| 89 | 0 | 20 | 0 | 0 |
| 91 | 0 | 20 | 0 | 0 |
| 98 | 0 | 0 | 0 | 20 |

**Table S8.**

Fixed-taxon RELL pseudo-bootstraps best-ranked topology frequencies for 15 unrooted topologies.

| Topology | SERR+LG | SERR+EDM | MARL+LG | MARL+EDM |
|---|---|---|---|---|
| 1 | 0 | 0 | 0 | 0 |
| 2 | 19 | 2 | 0 | 3 |
| 3 | 0 | 0 | 0 | 0 |
| 4 | 81 | 98 | 60 | 96 |
| 5 | 0 | 0 | 35 | 1 |
| 6 | 0 | 0 | 5 | 0 |
| 7 | 0 | 0 | 0 | 0 |
| 8 | 0 | 0 | 0 | 0 |
| 9 | 0 | 0 | 0 | 0 |
| 10 | 0 | 0 | 0 | 0 |
| 11 | 0 | 0 | 0 | 0 |
| 12 | 0 | 0 | 0 | 0 |
| 13 | 0 | 0 | 0 | 0 |
| 14 | 0 | 0 | 0 | 0 |
| 15 | 0 | 0 | 0 | 0 |

**Table S9.**

Compositional diversity model adequacy tests on the fixed-taxon data matrices. Data simulated under the EDM+G4 model are closest in amino acid compositional diversity to the empirical data. Bold values correspond to the simulated data that most closely resembles the empirical data.

| **Model** | **Amino acid diversity** | | **Z-score** |
|---|---|---|---|
| | **Empirical data** | **Simulated data** | |
| **MARL** | | | |
| LG+G4 | 5.1592 | 5.4859±0.005859 | 55.7581 |
| EDM+G4 | 5.1592 | 5.3082±0.005771 | **25.8182** |
| **SERR** | | | |
| LG+G4 | 6.2950 | 6.5769±0.007286 | 38.6836 |
| EDM+G4 | 6.2950 | 6.1735±0.006021 | **-20.1752** |

**Table S10.**

Randomised taxon-jackknife best-ranked topology frequencies for 15 unrooted topologies.

| **Topology** | **SERR+LG** | **SERR+EDM** | **MARL+LG** | **MARL+EDM** |
|---|---|---|---|---|
| 1 | 2 | 0 | 4 | 0 |
| 2 | 23 | 11 | 0 | 2 |
| 3 | 0 | 0 | 0 | 2 |
| 4 | 38 | 61 | 43 | 92 |
| 5 | 0 | 6 | 26 | 1 |
| 6 | 0 | 0 | 12 | 0 |
| 7 | 0 | 0 | 0 | 0 |
| 8 | 30 | 0 | 0 | 1 |
| 9 | 5 | 1 | 15 | 1 |
| 10 | 0 | 0 | 0 | 0 |
| 11 | 0 | 0 | 0 | 0 |
| 12 | 0 | 0 | 0 | 0 |
| 13 | 0 | 0 | 0 | 0 |
| 14 | 2 | 20 | 0 | 1 |
| 15 | 0 | 1 | 0 | 0 |

**Table S11.**

Simulations on 5-taxon Spiralia star-tree best-ranked topology frequencies for 15 unrooted topologies. For columns SERR+EDM and MARL+EDM, proportions are based on 10 alignments.

| Topology | SERR+LG EDM sim | SERR+EDM | MARL+LG EDM sim | MARL+EDM |
|---|---|---|---|---|
| 1 | 0 | 0 | 1 | 0 |
| 2 | 73 | 20 | 0 | 0 |
| 3 | 0 | 0 | 0 | 0 |
| 4 | 0 | 0 | 0 | 0 |
| 5 | 0 | 40 | 0 | 80 |
| 6 | 0 | 0 | 99 | 20 |
| 7 | 0 | 0 | 0 | 0 |
| 8 | 22 | 0 | 0 | 0 |
| 9 | 0 | 0 | 0 | 0 |
| 10 | 0 | 0 | 0 | 0 |
| 11 | 0 | 0 | 0 | 0 |
| 12 | 0 | 20 | 0 | 0 |
| 13 | 0 | 0 | 0 | 0 |
| 14 | 0 | 0 | 0 | 0 |
| 15 | 5 | 20 | 0 | 0 |
